# *Notch1* mutation drives clonal expansion in normal esophageal epithelium but impairs tumor growth

**DOI:** 10.1101/2021.06.18.448956

**Authors:** Emilie Abby, Stefan C Dentro, Michael W J Hall, Joanna C Fowler, Swee Hoe Ong, Roshan Sood, Christian W Siebel, Moritz Gerstung, Benjamin A Hall, Philip H Jones

**Affiliations:** Wellcome Sanger Institute, Hinxton CB10 1SA, UK; European Molecular Biology Laboratory, European Bioinformatics Institute, Cambridge CB10 1SD, UK; MRC Cancer Unit, University of Cambridge, Hutchison-MRC Research Centre, Cambridge Biomedical Campus, Cambridge CB2 0XZ, UK; Department of Discovery Oncology, Genentech, South San Francisco, CA, USA; Department of Medical Physics and Biomedical Engineering, University College London, Gower Street, London WC1E 6BT, UK

## Abstract

*NOTCH1* mutant clones occupy the majority of normal human esophagus by middle age, but are comparatively rare in esophageal cancers, suggesting *NOTCH1* mutations may promote clonal expansion but impede carcinogenesis^1–3^. Here we test this hypothesis. Visualizing and sequencing *NOTCH1* mutant clones in aging normal human esophagus reveals frequent biallelic mutations that block NOTCH1 signaling. In mouse esophagus, heterozygous *Notch1* mutation confers a competitive advantage over wild type cells, an effect enhanced by loss of the second allele. *Notch1* loss alters transcription but has minimal effects on epithelial structure and cell dynamics. In a carcinogenesis model, *Notch1* mutations were less prevalent in tumors than normal epithelium. Deletion of *Notch1* reduced tumor growth, an effect recapitulated by anti-NOTCH1 antibody treatment. We conclude that *Notch1* mutations in normal epithelium are beneficial as wild type *Notch1* promotes tumor expansion. NOTCH1 blockade may have therapeutic potential in preventing esophageal squamous cancer.

## Main text

As our tissues age they accumulate numerous somatic mutations^1,2,4,5^. Disruption of some genes confers a competitive advantage to mutant progenitor cells, which may go on to form mutant clones that spread through normal tissue. Such clonal expansions are often associated with genes recurrently mutated in cancer and are considered to be a first step in malignant transformation^2^. However, in the esophagus, the under-representation of *NOTCH1* mutants in esophageal cancer compared with normal aging epithelium suggests *NOTCH1* mutations may even protect against malignant transformation.

The NOTCH pathway regulates cell fate in development, tissue homeostasis and cancer^6,7^. NOTCH1 is a cell surface receptor composed of a ligand-binding extracellular domain (NEC) and a transmembrane and cytoplasmic subunit (NTM), interacting non-covalently through the negative regulatory region (NRR) (**Extended data Fig. 1a**). The NRR is composed of three Lin12-Notch repeats (LNR) and a heterodimerization domain (HD) that inhibits receptor activation in the absence of a ligand^8^. Binding of ligands expressed on adjacent cells to conserved epidermal growth factor (EGF) repeats in the NEC results in proteolytic cleavage events that release the cytoplasmic domain (NICD), which then migrates to the nucleus and alters the expression of target genes^8^. In the esophagus NOTCH1 protein is expressed in the proliferating cell compartment and regulates both development and adult tissue maintenance (**Extended data Fig. 1a,b**)^9^.

The role(s) of *NOTCH1* in esophageal carcinogenesis are controversial with different studies arguing *NOTCH1* is a tumor suppressor or promotes carcinogenesis^1,2,10–12^. Here we investigate how *NOTCH1* mutants take over the epithelium, the impact this has on transformation and whether NOTCH1 has potential as a therapeutic target in esophageal carcinogenesis^1,2^.

### *NOTCH1* mutant clones in human esophagus

Deep targeted sequencing studies have revealed numerous *NOTCH1* mutants in human esophagus but are unable to visualize clones within the tissue and resolve which *NOTCH1* mutation(s) or copy number alterations occur within each clone ^1^. To address these issues, histological sections of normal epithelium from middle-age and elderly donors were immunostained for NOTCH1 revealing continuous regions of different intensity, alternating with negative areas (**Fig. 1a**). To determine whether this staining pattern reflected *NOTCH1* mutations, we immunostained histological sections from 3 donors, aged 57, 68 and 76 years for NOTCH1 micro-dissected positive or negative areas and performed targeted sequencing for 322 genes associated with cancer (**Fig. 1b**). 176 protein altering somatic mutations were identified across 60 samples. Consistent with previous studies, the predominant mutations were in *NOTCH1*, *TP53* and *NOTCH2.* Protein altering mutations in *NOTCH1* represented a third of all mutation calls ^1,2^ (**Supplementary tables 1, 2, Supplementary Note**). *NOTCH1* mutations were near clonal with an average variant allele frequency (VAF) of 0.42 (**Fig. 1c**). 94% (15/16) of negative staining areas in the two oldest donors carried nonsense or essential splice mutations or indels in *NOTCH1* with copy neutral loss of heterozygosity (CNLOH) of the *NOTCH1* locus (human GRCh37-chr9:139,388,896-139,440,238) or a further mutation, likely to disrupt the second *NOTCH1* allele ^1^. Of the positive staining samples, 45% (20/44) carried a missense *NOTCH1* mutation and most of these had either CNLOH or a second mutation (**Fig. 1c, Extended data Fig. 1c-d, Supplementary Table 2**). To test if the mutations disrupted NOTCH1 function we stained consecutive sections from additional donors aged 22 to 77 years old for NOTCH1 protein and NICD1, which is detectable in the nucleus during active signaling with an anti-NICD1 antibody targeting a neo-epitope revealed after cleavage **(Fig. 1d, e, Extended data Fig. 1a, Supplementary table 3**). The proportion of epithelium with active NOTCH1 decreased with age (Kendall’s tau-b=-0.67, p=0.014). In older donors in whom *NOTCH1* mutations are common, we confirmed that NOTCH1 negative areas were associated with NICD1 loss. We also found occasional NOTCH1 positive NICD1 negative areas, consistent with the presence of missense mutant proteins that reach the cell membrane but have no detectable signaling activity (**Fig. 1d, e**). NICD1 positive and negative areas were histologically undistinguishable, with no significant differences in tissue thickness, cell density or the expression of the proliferation marker MKI67 (**Extended data Fig. 1e-h**). We conclude that the majority of *NOTCH1* mutant clones in aging human esophagus carry biallelic alterations that disrupt protein expression and/or function.

**Figure 1:**
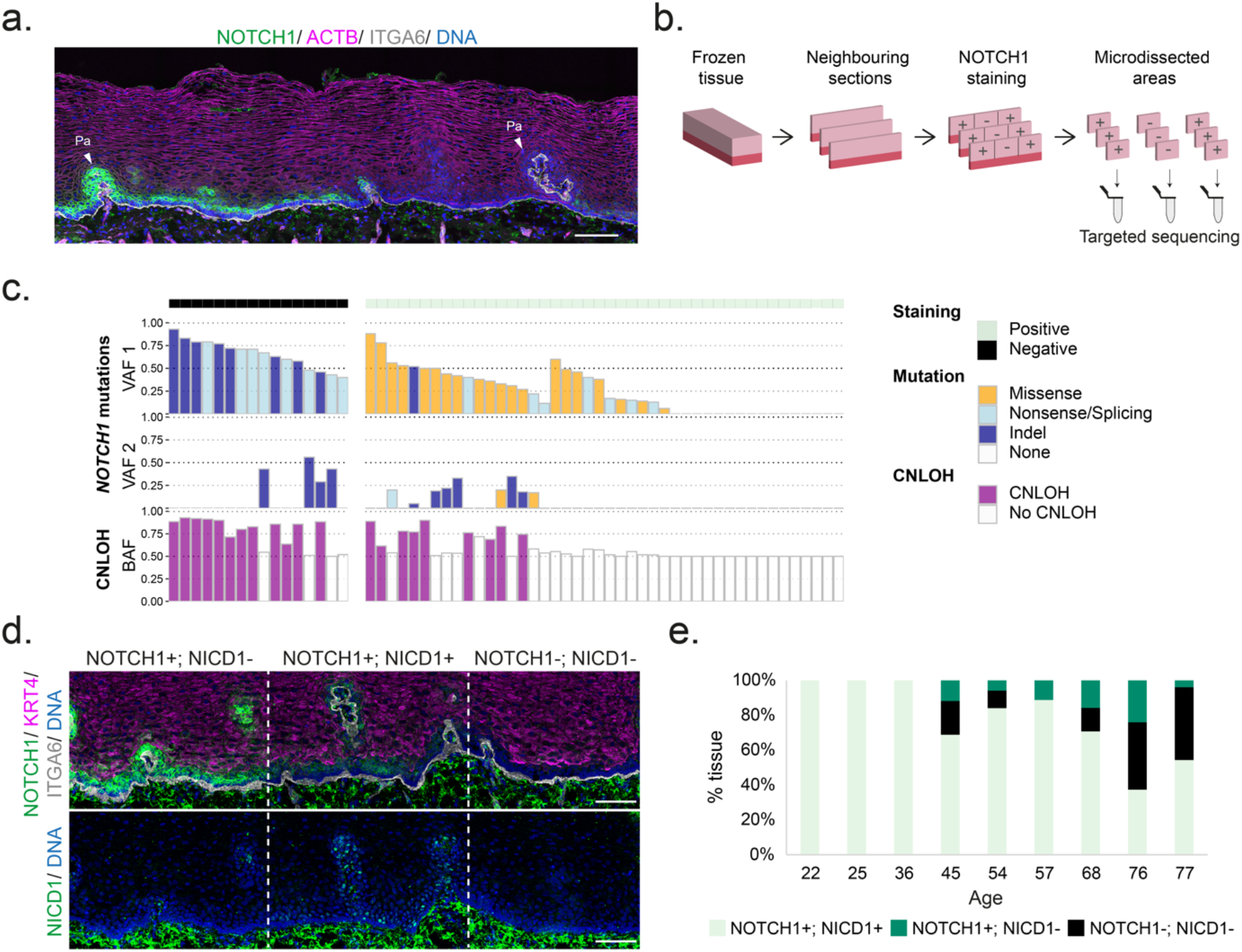
*NOTCH1* mutant clones in human esophageal epithelium. **a.** NOTCH1 (green) is expressed in the basal cell layer marked with ITGA6 (grey) but expression is lost in adjoining regions of esophagus. F-actin is magenta. Pa, papillae. Image is representative of sections from 3 elderly donors. Scale bar, 100µm. **b.** Protocol for **c**. Frozen esophagus from three donors were sectioned and stained for NOTCH1. Contiguous NOTCH1 positive (+) and negative (-) staining areas from successive sections were micro-dissected and processed for targeted sequencing. **c.** Results from **b**, showing NOTCH1 staining, *NOTCH1* mutation calling and copy number calling on chromosome 9. CNLOH, copy neutral loss of heterozygosity (n=60 samples from 3 donors aged 57, 68 and 76 years). **d.** Representative areas stained for NOTCH1 and NICD1 on successive sections of esophageal epithelium from an aged donor. Picture is representative of 6 middle-aged and elderly donors. Scale bar, 100µm. **e.** Proportion of tissue stained for NOTCH1 and associated NICD1 quantified in patients aged 22 to 77 years (total section length 4774-17988 µm per donor, n=9 donors). NOTCH1+; NICD1+ areas decrease significantly with age (Kendall’s tau-b correlation=-0.67, p=0.014). See **Supplementary tables 1-3**.

### *Notch1* mutation increases clonal fitness

To investigate how *NOTCH1* mutant clones colonize normal epithelium we tracked the fate of *Notch1* mutant clones in transgenic mice using lineage tracing. Mouse esophageal epithelium consists of layers of keratinocytes, with progenitor cells residing in the basal cell layer (**Extended data Fig. 2a**). Dividing progenitors generate either two progenitor daughters that remain in the basal layer, two differentiating daughters or one cell of each type. Differentiating cells exit the cell cycle, exit the basal layer and migrate towards the epithelial surface where they are shed. In wild type tissue, the probabilities of each progenitor outcome are balanced, generating equal proportions of progenitor and differentiated cells, maintaining cellular homeostasis across the progenitor population **(Extended data Fig. 2a)**^13,14^. Mutations can alter the balance of progenitor fate leading to excessive production of progenitors driving the growth of mutant clones^15,16^.

To follow the behavior of *Notch1* mutant clones we generated *AhCre^ERT^ Rosa26^floxedYFP^ Notch1^flox^ triple transgenic (YFPCreNotch1)* mice, a conditional *Notch1* model crossed with a genetic labeling system. A drug inducible form of *Cre* recombinase (*AhCre^ERT^*) was used to delete one or both conditional *Notch1* alleles in *Notch1^flox/wt^* or *Notch1^flox/flox^* animals and induce a separate conditional yellow fluorescent protein reporter allele (*Rosa26^floxedYFP^*) so YFP was expressed in recombined epithelial cells and their progeny (**Extended data Fig. 2b,c**)^13,17^. This model could be induced at low dose to allow recombination of in scattered single esophageal basal cells (clonal induction) or at a high level so a larger proportion of the esophageal cells was recombined (high induction) (**Extended data Fig. 2c,d**).

Excision of the *Notch1* allele and expression of the YFP reporter at the *Rosa26* locus can occur in in combination or separately, resulting in four possible populations of cells, N*otch1* mutant or wild type and YFP + or - (**Extended data Fig. 2c,d**). To identify induced *Notch1* mutant clones, we first analyzed the gene expression in wild type, *Notch1^+/-^* and *Notch1^-/-^* esophageal cells finding both *Notch1* mRNA and protein expression were halved in *Notch1^+/-^* compared with wild type keratinocytes and were abolished in *Notch1^-/-^* cells (**Fig.2 a-c, Extended data 3a-e**).

**Figure 2:**
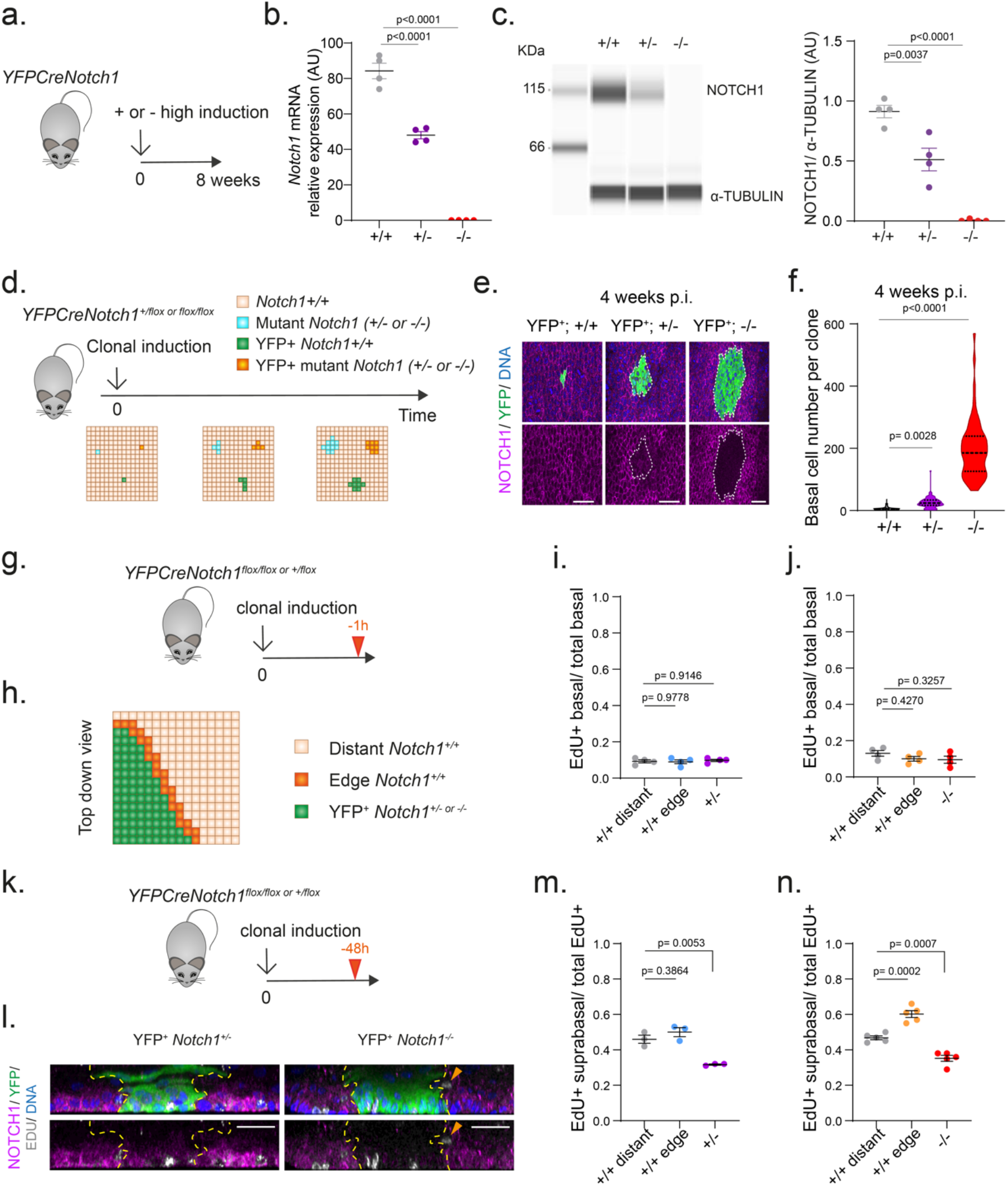
Lineage tracing of *Notch1* mutant clones. **a.** Protocol for **b** and **c**. *YFPCreNotch1^flox/flox^* or *^+/flox^* mice were induced to give a high level of recombination and aged for 8 weeks allowing *Notch1* mutant cells to colonize the epithelium. Non- induced *YFPCreNotch1^flox/flox^* mice were used as wild type controls (+/+). Esophageal epithelium was then collected. **b**. *Notch1* mRNA expression level measured by qRT-PCR (Mean ± SEM, each dot represents a mouse, n=4 mice). One-way ANOVA; adjusted p values from Tukey’s multiple comparisons test against wild type. **c.** Immune Capillary Electrophoresis assays for NOTCH1 transmembrane/intracellular domain (NTM1 + NICD1) and α-TUBULIN protein in esophageal epithelium (Mean ± SEM, each dot represents a mouse, n=4 mice). One-way ANOVA; adjusted p values from Tukey’s multiple comparisons test against wild type. **d.** *YFPCreNotch1^+/+^, YFPCreNotch1^+/flox^* and *^flox/flox^* mice were induced at clonal density. YFP+ wild type and YFP+ *Notch1* mutant clones (+/- or -/-) were analyzed at several time points (**Extended data Fig. 2c** and **3**). **e.** Projected view of 4 weeks post-induction (p.i.) wild type, *Notch1^+/-^* and *Notch1^-/-^* clones stained with NOTCH1 and YFP. White dashed lines delineate mutant clones. Scale bars: 30µm. **f.** Violin plots depicting number of basal cells per clone at 4 weeks post-induction (p.i.). Lines show median and quartiles. (n=143 clones from 3 +/+ mice, n=97 clones from 4 +/flox mice, n=63 clones from 9 flox/flox mice). One-way ANOVA; adjusted p values from Tukey’s multiple comparisons test against +/+. **g.** Protocol for short term clonal lineage tracing. *YFPCreNotch1^+/flox^* and *^flox/flox^* mice were induced at clonal level and injected with EdU 1h before collection. **h.** EdU+ cells were counted within the mutant YFP+ area (green), the wild type area at the edge of the mutant clone (orange) or distant from the clone (beige). **i, j**. Basal EdU positive and total basal cells were counted in *Notch1^+/-^* (**i**) or *Notch^-/-^* (**j**) mutant clones, in wild type cells at clone edges or distant from clones. Ratio of EdU positive basal cells to total number of basal cells was calculated for each cell population (Mean ± SEM, each dot represents a mouse; **i**, n= 4830/ 1584/ 4607 cells in +/+ distant/ +/+ edge/ +/- clones from 4 mice; **j**, n= 3967/ 1036/ 4279 cells in +/+ distant/ +/+ edge/ - /- clones from 4 mice). One-way ANOVA; adjusted p values from Tukey’s multiple comparisons test against +/+ distant. **k**. Protocol for short term clonal lineage tracing. Mice were induced at clonal level and injected with EdU 48h before collection. **l.** Side view of projected confocal z stacks of a *Notch1*^+/-^ clone at 3 months p.i. (left) and a *Notch1^-/-^* clone at 1 month (p.i. right) from animals injected with EdU 48h before collection. NOTCH1 (magenta); YFP (green); EdU (grey); DNA (blue). Yellow dashed lines show the edges of the clones. Orange arrow points to a differentiating cell adjacent to a clone. Images are representative of clones analyzed in respectively 3 *Notch1*^+/-^ and 5 *Notch1^-/-^* mice. Scale bars: 30µm. **m, n.** EdU was injected as in **k**. EdU positive cells were counted in *Notch1^+/-^* (**m**) or *Notch1^-/-^* (**n**) mutant clones, in wild type cells at the edge (+/+ edge) or distant from (+/+ distant) clones. Ratio of EdU positive suprabasal cells was calculated for each cell population. (Mean ± SEM, each dot represents a mouse; **m**, n= 471/ 300/ 525 EdU+ cells in +/+ distant/ +/+ edge/ +/- clones from 3 mice; **n**, n=1304/ 723/ 1318 EdU+ cells in +/+ distant/ +/+ edge/ -/- clones from 5 mice). One-way ANOVA; adjusted p values, Tukey’s multiple comparisons test against +/+ distant. AU, arbitrary unit.

We then performed genetic lineage tracing by inducing recombination in scattered single progenitor cells in control (*Notch1^+/+^*), heterozygous (*Notch1^+/ flox^*) and homozygous (*Notch1 ^flox / flox^*) mutant animals. The growth of the YFP expressing clones in each mouse was detected by imaging sheets of epithelium stained for both YFP and NOTCH1 at multiple time points. We were able to identify YFP+ *Notch1^+/-^* and YFP+ *Notch1^-/-^* clones from the intensity of NOTCH1 immunostaining (**Fig. 2d,e, Extended data Fig. 3f-i**). The number and location of cells in YFP expressing clones of each genotype was then scored across the time course using 3D confocal imaging.

The size of YFP+ *Notch1^+/-^* clones was significantly increased compared with those of wild type YFP+ *Notch1^+/+^* clones at all time points. YFP+ *Notch^-/-^* clones were larger still (**Fig. 2e,f, Extended data Fig. 3j-k**). To examine the cellular mechanisms underlying mutant clonal expansion we used short term tracking by labelling cycling cells with the S phase probe EdU, and subsequently imaging the distribution of their EdU positive progeny. One hour after labelling the proportion of EdU positive cells was similar in wild type, *Notch1^+/-^* and *Notch1^-/-^* cells, arguing the rate of cell division was similar for progenitors of all three genotypes (**Fig. 2g-j**). At 48 hours after labelling, the proportion of suprabasal EdU positive cells reflects the rate at which basal cells differentiate and stratify out of the basal cell layer. This was significantly decreased within YFP+ *Notch1^+/-^* and YFP+ *Notch1^-/-^* clones compared with wild type cells distant from the clone. However, in *wild type* cells immediately adjacent to YFP+ *Notch1^-/-^* cells the proportion of EdU positive suprabasal cells was significantly increased, consistent with previous reports that Notch-inhibited keratinocytes promote the differentiation of adjacent wild type cells (**Fig. 2k-n**) ^16,18^.

These observations and constraints were integrated into a simple Wright-Fisher style quantitative model of ‘neighbor constraint’, in which competition is driven by local fitness differences between cells. Mutant clones expand until they collide with other mutant clones of similar fitness, at which point they revert to neutral competition ^19^. We fitted this model to the clone size data to estimate the fitness of mutant clones relative to wild type cells with a fitness of 1. The inferred fitness for *Notch1^+/-^* clones was substantially greater than that of wild type cells and the inferred fitness of *Notch1^-/-^* clones markedly higher than that of heterozygous clones (**Extended data Fig. 4a-d, Video 1, Supplementary Note**). We conclude that loss of one allele of *Notch1* is sufficient to drive clonal expansion, but the competitive fitness of clones is substantially augmented when the second allele is lost.

### *Notch1* haploinsufficiency is pivotal in the colonization of normal epithelium

Clones generated by the transgenic deletion of *Notch1* alleles may not reflect the behavior of the sporadic *Notch1* mutant clones carrying truncating, missense or indel mutations that develop during aging. We therefore investigated the appearance of spontaneous *Notch1* mutant clones in control *YFPCreNotch1^+/+^* mice, comparing this with highly induced YFPCre*Notch1^+/-^* animals which we used to model the effects of loss of a second *Notch1* allele within heterozygous epithelium. Both strains were aged for over a year prior to immunostaining the epithelium for NOTCH1 (**Fig. 3a**). We observed a progressive increase in the area of epithelium staining negative for NOTCH1 protein in both induced strains, rising to 12 % of *Notch1 ^+/+^* and 78% of *Notch1 ^+/-^* epithelium by 65 weeks (**Fig. 3b,c**). Widespread loss of NICD1 staining was also observed in the aged *Notch1^+/-^* tissue (**Extended data Fig. 5a,b**). These findings show that esophageal epithelium is colonized by *Notch1* mutant clones during aging, as is the case in humans, and that loss of the remaining *Notch1* allele in *Notch1^+/-^* epithelium increases competitive fitness.

**Figure 3:**
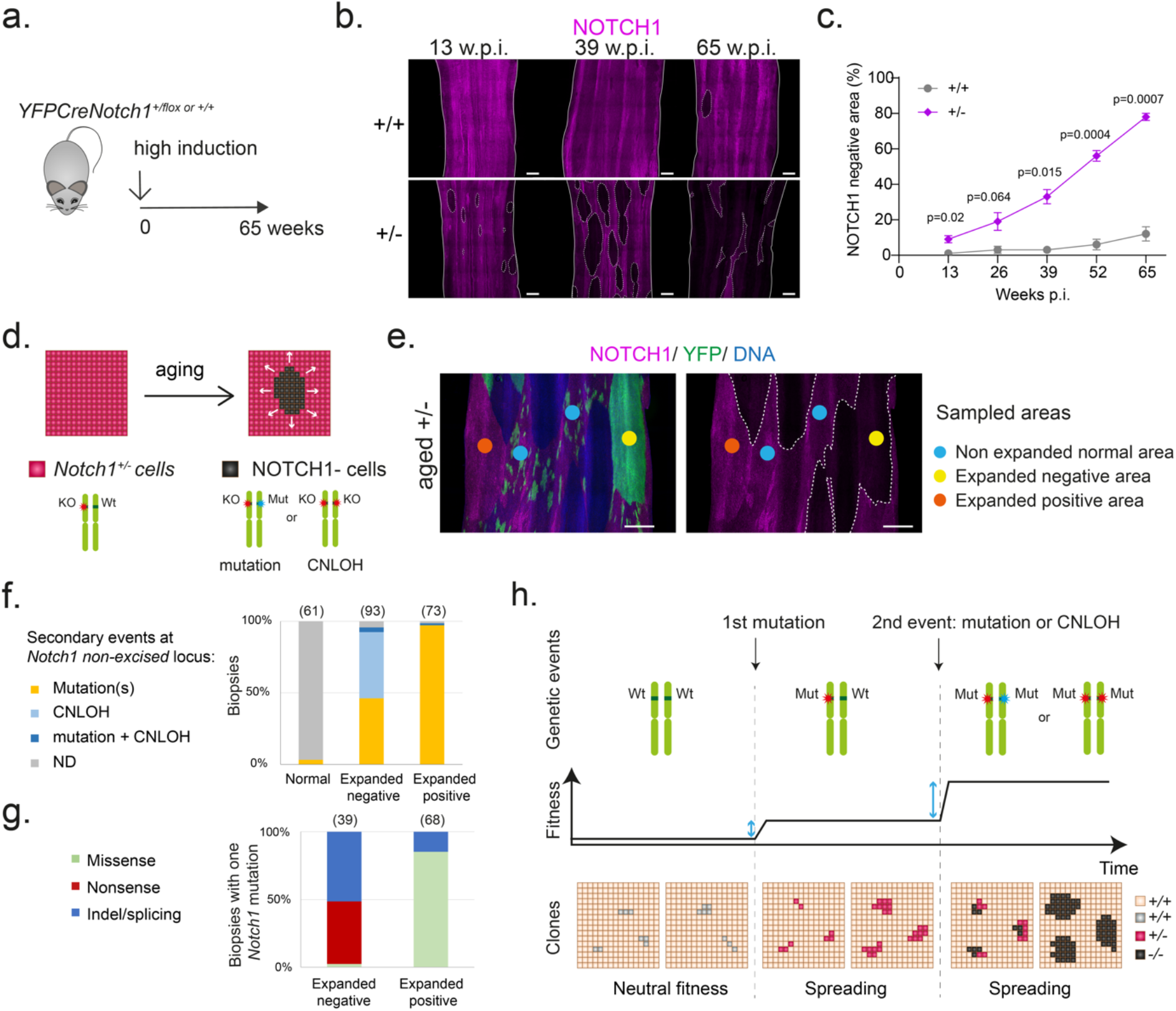
Notch1 colonizes aging esophageal epithelium. **a.** *YFPCreNotch1^+/flox^* and *^+/+^* mice were induced at high level and aged for 65 weeks. **b.** Representative NOTCH1 staining in esophageal epithelium of *Notch1^+/+^* and *Notch1^+/-^* mice at the indicated time points. White dashed lines delineate negative areas, solid lines delineate tissue edges. Images are representative confocal z stack of entire epithelium from 3 mice per time point. W.p.i, weeks- post-induction. **c.** Percentage of NOTCH1 negative area increases with age in *Notch1^+/+^* (Kendall’s tau-b correlation=0.56, p=0.0062) and *Notch1^+/-^* (Kendall’s tau-b correlation=0.91, p=8.3e-6) esophagi (Mean± SEM, n=3 mice per time point). P values shown are from two-sided Welch’s t-test. Scale bars: 500µm. **d**. Schematic of *Notch1^+/-^* cells (purple cells) showing the spontaneous appearance of expanding NOTCH1 negative cells (black) with aging, possibly caused by genetic events affecting the *Notch1* locus. **e.** Induced *YFPCreNotch1^+/-^* mice were aged 54 to 78 weeks-old, when esophageal epithelium was collected and stained for NOTCH1 (magenta) and YFP (green). Expanding areas devoid or fully stained with YFP appeared distinct from normal appearing areas marked with a patchwork of small YFP+ clones. Expanded NOTCH1 negative (yellow) and positive (orange) areas and normal appearing areas (blue) were isolated for targeted sequencing (n=246 biopsies from 10 mice). Colored circles show the sampled areas. White dashed lines delineate negative areas. Scale bar: 500µm. **f.** Proportion of normal appearing, expanded NOTCH1 negative and expanded NOTCH1 positive biopsies with *Notch1* mutations or copy number alternations. **g**. Proportion of NOTCH1 negative and positive areas carrying a secondary missense, nonsense or indel/splicing *Notch1* mutation. For **f** and **g**, number of samples is shown in brackets, redundant samples, defined as biopsies sharing the same mutation and separated by <1mm were counted once. **h.** Summary scheme of colonization *by Notch1* clones. Clonal fitness increases from monoallelic and biallelic *Notch1* mutation resulting in a selective pressure (blue arrows) for biallelic gene alterations. P.i., post-induction, Wt, wild-type; KO, knock-out allele lacking *Notch1* exon 1; Mut, mutation., ND, none detected. See **Supplementary table 4**.

In order to isolate and sequence potential clones we used NOTCH1 staining in combination with the YFP reporter. Aging *Notch1^+/-^* epithelium contained multiple ovoid areas far larger than most YFP labelled clones. Some of these were positive for YFP (YFP+), whereas in others all YFP clones were absent (YFP-), suggesting they had been displaced by an expanding clone (**Fig. 3d,e**). These NOTCH1 positive and negative ‘expanded’ areas were imaged, dissected out and targeted sequencing of 73 Notch pathway and cancer related genes was performed on a total of 246 biopsies (**Supplementary tables 1 and 4)**. We analyzed CNLOH and all mutations with a VAF higher than 0.2, as below this threshold mutations were considered unlikely to drive the presumed clonal expansion. 97% (180/185) of the sequenced YFP+ and YFP- expanded areas had either *Notch1* protein-altering mutations with VAF≥0.2 or CNLOH at Chromosome 2 involving the *Notch1* locus (GRCm38 - chr2:26,457,903- 26,503,822). In contrast, only 2/61 samples from non-expanded areas carried *Notch1* mutations and none had CNLOH (**Fig. 3f, Extended data Fig. 5c, d, Supplementary table 4**). A small number of mutations in other genes were also called with VAF≥0.2, however they were either found coupled with a high VAF *Notch1* genetic alteration, so may represent passenger mutations within a *Notch1* mutant clone or were found in non-expanded areas. Expanded areas were thus associated with genetic events affecting *Notch1* locus. Furthermore, 94% (169/180) of the expanded areas with *Notch1* altering events carried only a single event (about 50% one *Notch1* protein-altering mutation and the remainder CNLOH) with an average VAF 0.44, consistent with them being clones carrying spontaneous genetic events affecting the non-recombined *Notch1* allele. (**Fig. 3e-g**, **Supplementary Table 4, Extended data Fig. 5c,d**). NOTCH1 expressing clones harbored missense mutations (87%) or Indels (13%) (**Fig. 3g**). Overall the effect of these mutations on NOTCH1 expression is consistent with our observations in human tissue (**Fig.1**).

To test the functional impact of missense mutations we used an *ex-vivo* assay of NOTCH1 function **Extended data Fig. 5e-j, Supplementary table 4**) ^20^. *YFPCreNotch1^+/-^* tissues analyzed in **Fig. 3e-g** were incubated with EDTA at 37°C prior to fixation. Depletion of Ca2+ using EDTA promotes the cleavage of NEC and NTM subunits without ligand binding, thought to be due to conformational changes in the LNR domains (**Extended data Fig.1a**)^20^. This results in the release and nuclear migration of NICD. Some NOTCH1 positive clones displayed NICD nuclear staining, but in others nuclear NICD was absent (**Extended data Fig, 5e**). Nuclear NICD staining clones were enriched in missense mutations located in the ligand binding site, EGF repeats 8-12, whereas non-nuclear staining clones were enriched in NRR domain mutants **(Extended data Fig. 5g, h** p=0.001, Chi-square test). Most of the mutated residues in the ligand binding domain had highly destabilizing properties, consistent with them impairing Notch signaling by disrupting ligand binding, a process bypassed in the ETDA assay (**Extended data Fig. 5i)**^21,22^. The NRR domain mutants were clustered in the LNR1 and 2 domains (**Extended data Fig. 5j)**^23^. This contrasts with *NOTCH1* activating mutations in human T cell acute lymphoblastic leukemia (T- ALL), collated from COSMIC (https://cancer.sanger.ac.uk/cosmic), which cluster in the HD domain of the NRR and promote the cleavage of NEC without ligand interaction (two-sided Fisher exact test comparing mutation counts in the LNR1-2 and LNR3-HD sub-regions of the NRR, p=1.48e^-10^ **Extended data Fig. 5j)** ^24,25^. This observation associated with the absence of detectable nuclear NOTCH1 after EDTA treatment in this group of clones suggests that the mutations they carry at the NRR domain may lock it into its auto-inhibited conformation, preventing the cleavage of NOTCH1.

We conclude that most NOTCH1 expressing clones carry mutations disrupting NOTCH1 function. Overall, the emergence of large clones carrying mutations or CNLOH in *Notch1* heterozygous mice is consistent with spontaneous mutation of the remaining *Notch1* allele conferring a fitness advantage over neighboring heterozygous cells.

Collectively these observations reveal that haploinsufficiency is key for the normal esophagus to be colonized so effectively by *Notch1* mutants. In simulations, neutral mutations do not spread across the tissue and are likely to disappear from the tissue over time. However, loss of one allele biases mutant progenitor cell fate towards the production of progenitors, increasing the likelihood that mutant clones will expand and persist in the epithelium (**Extended data Fig. 4e, f, Video 2**) *Notch1* inactivated cells have a further fitness advantage, so that subclonal loss of the second allele within a persisting heterozygous clone will generate cells that outcompete both *Notch1^+/+^* and *Notch1^+/-^* neighbors (**Fig. 3h)**. Over time biallelic clones will thus populate the tissue. This model explains the high prevalence of clones with *NOTCH1* mutation and CNLOH in aging human esophagus.

### *Notch1* loss alters transcription but not tissue function

The replacement of large areas of the esophageal epithelium by clones lacking functional NOTCH1 might be expected to have cellular and tissue level phenotypes. To explore the effects of *Notch1* loss we first performed bulk RNA sequencing on peeled esophageal epithelium from *Notch1* wild type, heterozygous and homozygous mice (**Fig. 4a-e**). *YFPCreNotch1^+/flox^* and *YFPCreNotch1^flox/flox^* mice were induced at a high level and aged for 8 weeks so that the recombined cells occupied almost all the esophageal epithelium (**Fig. 4a, Extended data Fig.3e, Fig.2-c**). RNA-seq was performed on the peeled esophageal epithelium from these mice and wild type littermates. In comparison with wild type tissue, 20 genes in *Notch1^+/-^* and 227 genes *Notch1^-/-^* esophagus were differentially expressed **(Fig. 4c-e, Supplementary table 5**). These included previously identified *Notch1* regulated genes, *Igfbp3* and *Sox9*, but transcription of canonical *Notch* targets such as *Hes1* and *Nrarp* was not significantly altered (**Supplementary table 5**) ^16,26,27^. Gene Ontology analysis showed genes regulating biological processes related to squamous epithelia were not significantly affected (EnrichGO q-value > 0.05), though transcripts of genes involved in DNA replication were downregulated (q= 2.6x 10^-7^) (**Supplementary table 5**).

**Figure 4:**
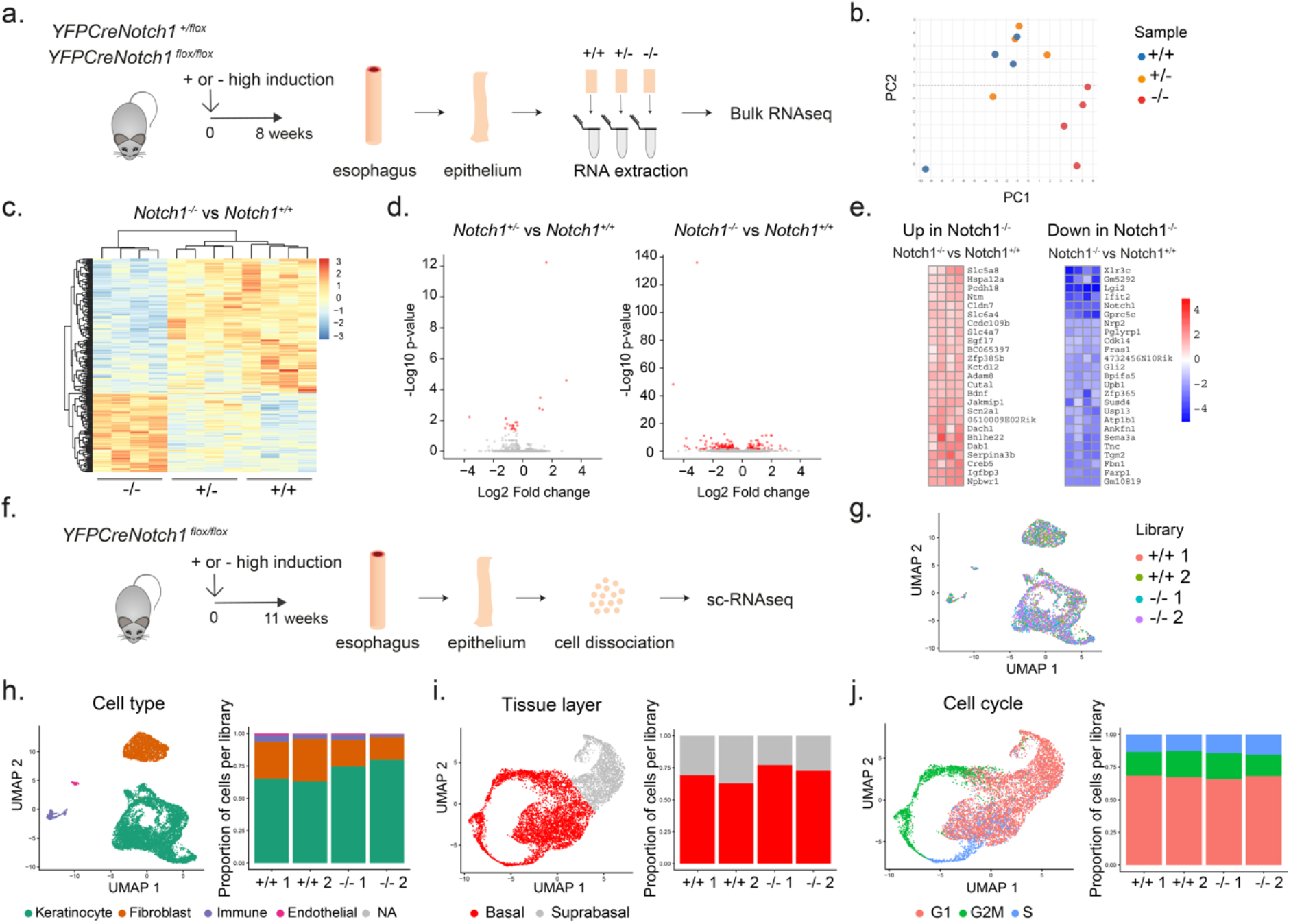
*Notch1* loss alters transcription but not tissue composition or cell dynamics. **a.** *YFPCreNotch1^flox/flox^*, *YFPCreNotch1^+/flox^* mice were highly induced and aged for 8 weeks to allow the mutant cells to cover the esophageal epithelium. Uninduced *YFPCreNotch1^flox/flox^* mice were used as wild type controls (+/+). Esophageal epithelium was collected and peeled and samples were processed for bulk RNAseq (n=4 mice per group). **b.** Principal component analysis (PCA) plot showing the four biological replicates of *Notch1^+/+^*, *Notch1^+/-^* and *Notch1^-/-^* esophageal epithelium samples in two dimensions. **c**. Hierarchical clustering and heat map showing differentially expressed genes between *Notch1^-/-^* and control tissues, in all three genotypes. **d**. Volcano plots showing *Notch1^+/-^* vs control (left) and *Notch1^-/-^* vs control (right) RNAseq analyses. Differentially expressed genes are indicated in red **e.** RNA-seq generated heat maps showing Log2 fold changes of the 25 top differentially expressed genes (DEG) in *Notch1^-/-^* compared to *Notch1^+/+^* tissues, Dseq2 analysis, adjusted p-value <0.05. **f.** *YFPCreNotch1^flox/flox^* mice were highly induced and aged for 11 weeks, allowing the mutant cells occupy the esophageal epithelium. Controls were uninduced *YFPCreNotch1^flox/flox^* mice (+/+). Esophageal epithelium was dissociated and sequenced. **g**. UMAP plot shows an overlay of 1500 cells from each library (n=2 mice per genotype; +/+1, n=2454; +/+2, n=3194; -/-1, n=1929; -/-2, n=5534). **h**. Left, UMAP plot showing cell types identified via sc-RNAseq. Right, stacked bar chart shows the proportion of cell types per library. NA, not available **i.** Left, UMAP plot of keratinocytes. Right, stacked bar chart shows the estimated proportion of keratinocytes per library belonging to the basal or suprabasal layers. **j.** Left, UMAP plot of keratinocytes and the likely cell cycle phase they belong to (G1, G2/M or S). Right, stacked bar chart showing proportion of keratinocytes per library assigned to each cell cycle phase. See **Supplementary tables 5 and 6.**

To define the effect of Notch1 loss on cell lineages and cellular states we performed single cell RNA sequencing (scRNA-seq). *YFPCreNotch1^flox/flox^* mice were induced at high density and aged so that the recombined keratinocytes covered the tissue entirely (**Fig. 4f, Extended data Fig.6a**). scRNA-seq was performed on dissociated epithelium from these mice and uninduced littermate controls **(Fig. 4f-j, Extended data Fig. 6b-e, Supplementary table 6**). After filtering out poor quality cells, a total of 13111 cells remained for analysis, from two biological replicates per genotype (**Fig. 4g, Supplementary Note**). We analyzed the cells using the Seurat software package, which identified cell clusters corresponding to four lineages **(Fig. 4h**, **Extended data Fig. 6c,e)** ^28^. The proportions of keratinocytes, fibroblasts immune and endothelial cells were similar in the control and *Notch1^-/-^* esophagus (**Fig. 4h**). We next applied the Seurat pipeline to the keratinocytes alone **(Extended data Fig. 6d**). The analysis revealed the existence of a continuum of keratinocytes states, from progenitors expressing *Krt14*, to differentiating cells expressing *Krt4* or *Tgm3* to terminally differentiated cornified cells expressing *Lor* (**Extended data Fig. 6e**). We used the expression level of these markers to discriminate basal and suprabasal cells on the UMAP plot (**Supplementary Note**). The analysis revealed similar proportions of keratinocytes in each differentiation state in the control and *Notch1^-/-^* tissue (**Fig. 4i**). Cell cycle gene expression patterns in keratinocytes also showed minimal differences between *Notch1^-/-^* and wild type tissues (**Fig. 4 j**). Overall, unbiased exploratory RNA-seq approaches did not evidence major alterations in tissue function or cellular composition due to loss of *Notch1* in esophageal epithelium. This conclusion was supported by imaging of the esophagus from induced and aged *Notch1^flox/flox^* mice and control littermates. Tissue thickness, basal cell density and immunostaining for differentiation markers KRT14, KRT4 and LOR and the proliferation marker Ki67 were similar in both genotypes, (**Fig.5a-d**).

**Figure 5:**
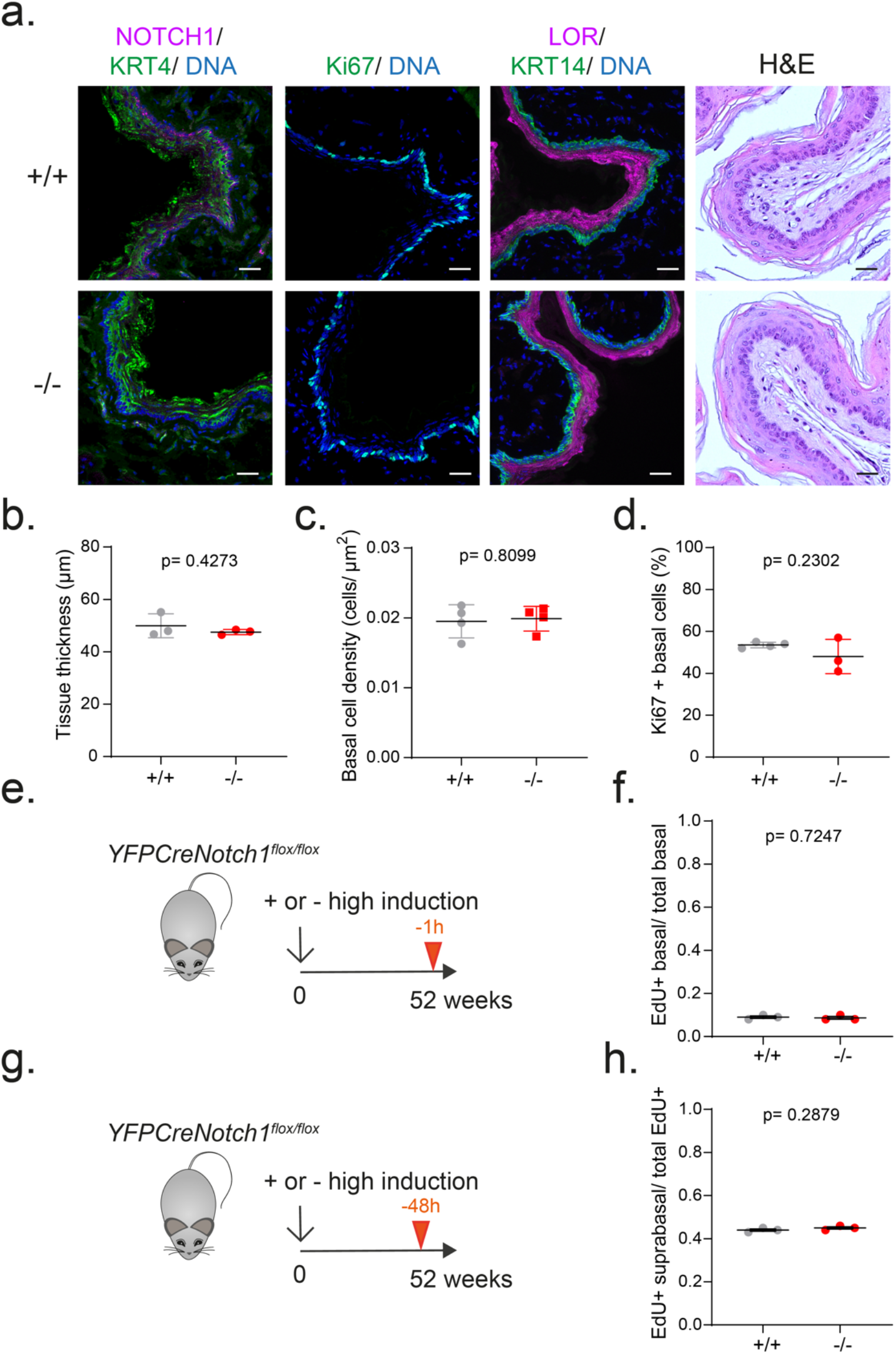
Differentiation and homeostasis in *Notch1* mutant mouse tissue. **a.** *Notch1^+/+^* and *Notch1^flox/flox^* mice were induced at high dose and aged as in (e) and (g) and tissue was collected. After sectioning, tissue was stained for basal cell marker KRT14, NOTCH1, proliferation marker Ki67, differentiation markers KRT4 and LOR and with Haematoxylin and eosin (H&E). Images are representative from 3 mice of each genotype. Scale bars, 30 μm. **b.** Thickness of the epithelium was measured on H&E scanned sections (Mean ± SEM, each dot represents a mouse, n=3 mice). Twotailed unpaired Student’s t-test. **c.** Epithelium basal cell density was measured on whole mount tissue. (Mean ± SEM, each dot represents a mouse, +/+, n=4097cells from 4 mice; -/-, n= 3964 cells from 4 mice). Two-tailed unpaired Student’s t-test. **d.** Proportion of proliferative basal cells were measured on sections stained for Ki67, KRT14 and DAPI. (Mean ± SEM, each dot represents a mouse, +/+, n= 1548 cells from 4 mice; -/-, n= 1129 cells from 3 mice). Two-tailed unpaired Student’s t-test. **e, f.** Highly induced or uninduced control *Notch1^flox/flox^* mice were aged for at least 52 weeks and injected with EdU 1h before collection (**e**). Ratio of EdU positive basal cells on total number of basal cells was calculated (**f**) (Mean ± SEM, each dot represents a mouse, +/+, n= 2754 cells from 3 mice; -/-, n= 2565 cells from 3 mice). Two-tailed, unpaired Student’s t-test. **g, h.** Highly induced or uninduced control *Notch1^flox/flox^* mice were aged for at least 52 weeks and injected with EdU 48h before collection (**g**). Ratio of EdU positive suprabasal cells. (Mean ± SEM, each dot represents a mouse, (+/+, n= 2687 EdU+ cells from 3 mice; -/-, n= 2201 EdU+ cells from 3 mice). Two-tailed unpaired Student’s t-test.

To augment these observations, we tracked cell dynamics by short term lineage tracing with EdU. 1h labelling indicated no significant difference in the proportion of S phase cells in *Notch1^-/-^* and control esophagus, despite the altered levels of DNA replication gene transcripts noted above (**Fig. 5e,f**). The proportion of suprabasal EdU positive cells 48 hours post labelling was also similar in both strains, an observation that contrasts with *Notch1^-/-^* cells in expanding clones. This suggested that once the esophagus has been fully colonized the behavior of *Notch1^-/-^* progenitor cells reverts towards normal, sustaining a normal appearing tissue (**Fig.5 g,h**) ^16,19^.

### *Notch1* loss alters tumor growth

*NOTCH1* mutations are under strong positive selection in normal human esophagus and occupy 30% to 80% of the epithelium as early as middle-age but are only found in about 10% of esophageal squamous cell carcinomas (ESCC) ^1,3^. This disparity hints at a role for wild type *NOTCH1* in malignant transformation. We therefore explored the role of *Notch1* in murine neoplasia. *Notch1* wild type mice were treated with the mutagen Diethylnitrosamine (DEN), and Sorafenib, a multikinase inhibitor that increases cell turnover and promotes the formation of high grade dysplastic lesions ^29^. After aging for 28 weeks, tissue was collected (**Fig. 6a**). Deep targeted sequencing of 73 cancer associated and Notch pathway genes was performed on dissected macroscopic tumors and a gridded array of normal epithelium (**Fig.6a**, **Supplementary table 7**).

**Figure 6:**
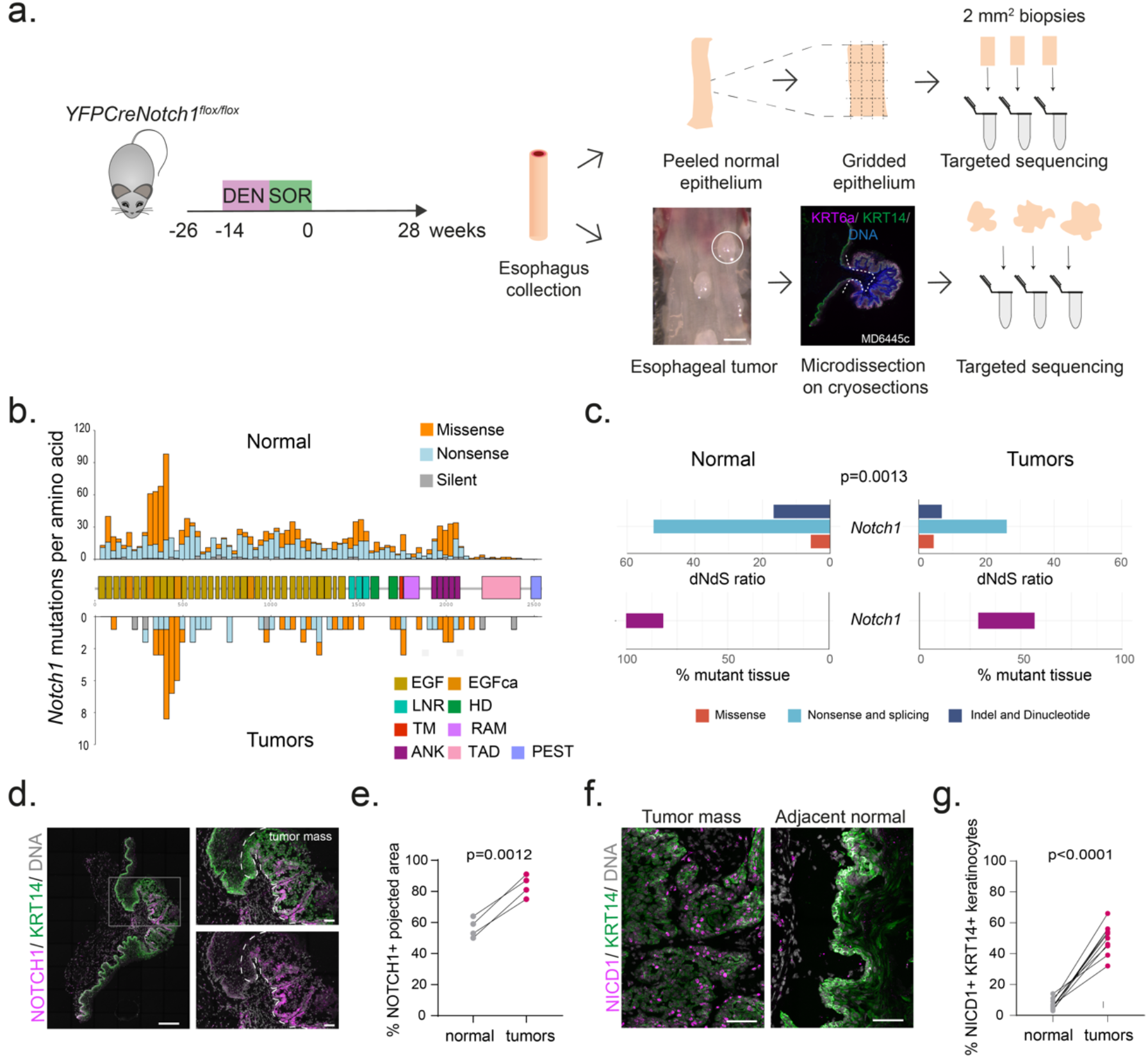
Tumors retain functional *Notch1* in carcinogenesis. **a.** Uninduced *YFPCreNotch1^flox/flox^* mice were treated with DEN and Sorafenib. Tissue was collected 28 weeks after treatment. Tumors were dissected from underlying submucosa and normal epithelium was cut into a gridded array of 2 mm^2^ samples prior to targeted sequencing. Scale bar, 1mm. **b.** Number of *Notch1* mutations per amino-acid is plotted by NOTCH1 protein domains in *Notch1^+/+^* normal gridded biopsies (upper) and *Notch1^+/+^* tumors (lower) (Normal, n=115 biopsies from 6 mice; Tumors, n=17 biopsies from 7 mice). **c**. dN/dS ratio for *Notch1* mutations and proportion of *Notch1* mutant tissue in normal epithelium (n=115 biopsies from 6 mice) and tumors (n=17 biopsies from 7 mice). Two-tailed p-value, likelihood ratio test of dN/dS ratios ^1^. **d.** Representative NOTCH1 (magenta) and KRT14 staining (green) in tumors and surrounding tissue, DNA is grey. Image typical of 10 tumors from 6 animals. White dashed line delineates tumor. Left scale bar, 250μm; Right scale bar, 50μm. **e.** Proportion of NOTCH1+ staining area in normal epithelium and tumors from the same control animals (each dot represents a mouse, n=40 tumors from 4 mice). Two-tailed paired Student’s t-test. **f.** Representative images showing nuclear NICD1 protein (magenta) in keratinocytes (KRT14, green) inside a tumor in comparison to the normal adjacent tissue. Image typical of 10 tumors from 6 animals. Scale bars, 50μm. **g.** Proportion of KRT14 positive keratinocytes with nuclear NICD1 staining in tumors and surrounding epithelium in the same sections (each dot represents a tumor, n=10 tumors from 6 mice). Two-tailed paired Student’s t-test. See **Supplementary table 7.**

The normal epithelium contained a high density of clones carrying protein altering mutations (**Supplementary table 7)**. To determine which genes conferred a clonal advantage, we calculated the ratio of silent to protein altering mutations in each gene, dN/dS ^5,30^. Mutant genes under positive selection with a dN/dS ratio significantly above 1 (q<0.05) were the Notch pathway genes *Notch1, Notch2* and *Adam10,* plus *Fat1, Trp53* and *Arid1a*, all of which are selected in normal human esophagus along with *Ripk4* and *Chuk* (**Supplementary table 7**) ^1,19^.

In tumors, the most selected gene was the known mouse esophageal tumor driver *Atp2a2*, which is rarely mutated and not selected in normal epithelium **(Extended data Fig. 7a,b. Supplementary table 7)** ^31,32^. *Notch1* mutations were similar to the protein disrupting mutants in normal tissue with no evidence of activating mutations. However, *Notch1* mutants were both less selected and less prevalent in tumors than in the adjacent tissue (**Fig. 6b, c**, **Extended data Fig. 7a,b, Supplementary table 7**). Immunostaining confirmed a higher proportion of cells staining positive for NOTCH1 and NICD1 in tumors than in the normal epithelium (**Fig. 6d-g**). These findings indicate *Notch1* wild type cells are more likely to contribute to tumors than those carrying *Notch1* mutants, consistent with the sequencing results in humans.

To test if wild type *Notch1* is required for tumor formation we used a high induction protocol to delete one or both alleles in the entire esophageal epithelium of *Notch1^flox/flox^* and *Notch1 ^+/flox^* mice prior to treatment with DEN and Sorafenib. Uninduced littermates were used as wild type controls (**Fig. 7a**). Detailed microscopic and macroscopic analysis of the tissue revealed that the density of tumors was similar in all three genotypes, arguing *Notch1* is not required for tumor formation **(Fig.7 b,c**). However, tumors were significantly smaller in *Notch1^-/-^*, compared to *Notch1^+/-^* and *Notch1^+/+^* animals (**Fig. 7d,e, Extended data Fig. 7c)**. NOTCH1 immunostaining confirmed the loss of *Notch1* expression in tumors from induced *Notch1^flox/flox^* mice. (**Extended data Fig. 7d**). Immunostaining for differentiation markers (LOR, KRT14) revealed that the tumors had similar staining for proliferation (Ki67), apoptosis (cleaved caspase 3), endothelial cell (CD31) and immune cell (CD45) markers in lesions from *Notch1^-/-^* and *Notch1^+/+^* esophagus (**Fig. 7e**, **Extended Data Fig. 7d-f)**. The cell-cell adhesion protein CDH1 (E-cadherin) is down regulated in human ESCC and loss of CDH1 signaling promotes tumorigenesis in mice ^33^. Interestingly, *Notch1* wild type tumors displayed focal loss of CDH1 expression, a feature that was not seen in the *Notch1^-/-^* tumors (**Fig.7 f,h**).

**Figure 7:**
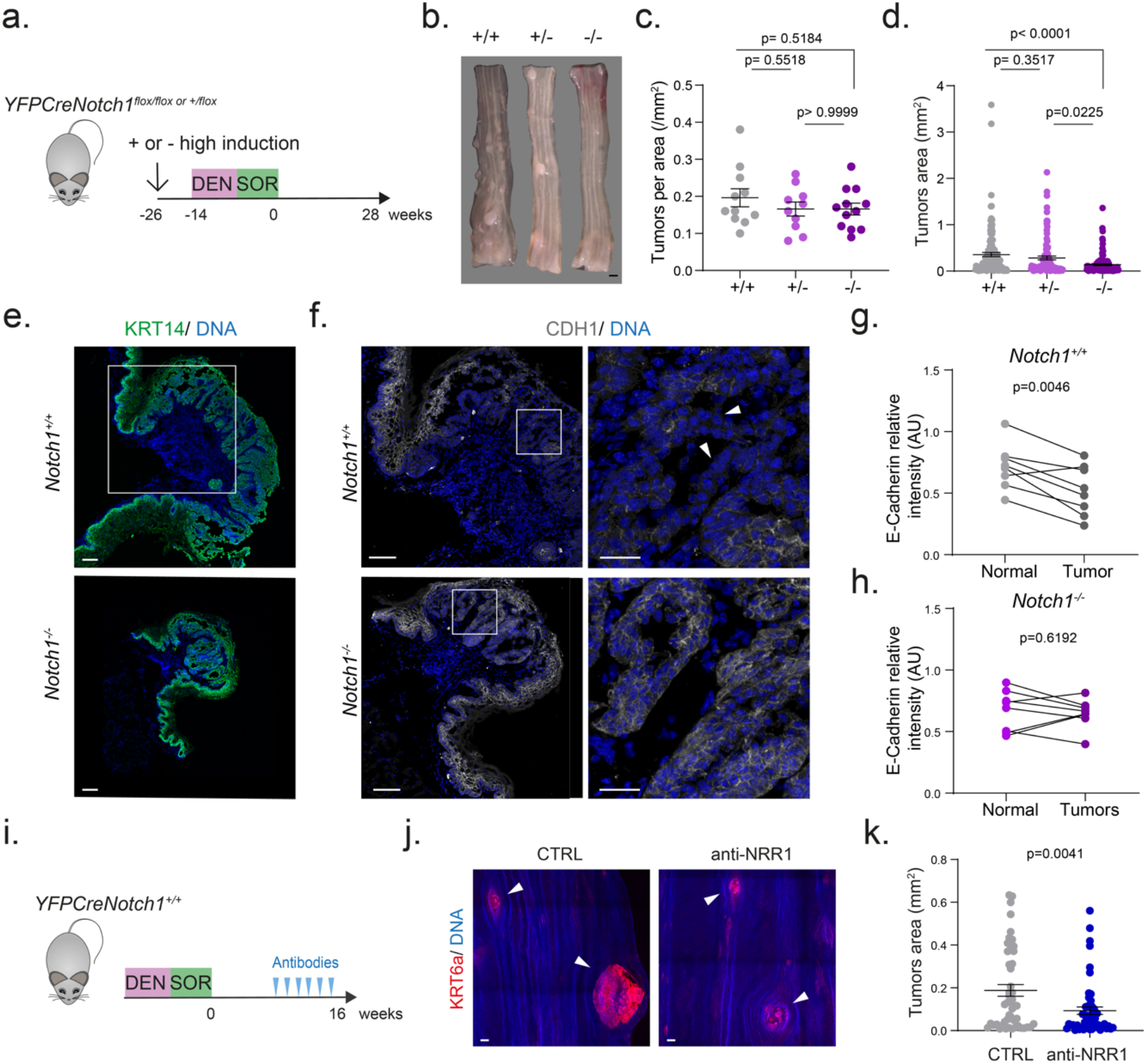
Tumor growth is reduced by *Notch1* inactivation. **a.** Highly induced *YFPCreNotch1^+/flox^* and *YFPCreNotch1^flox/flox^* mice or uninduced control mice were treated with DEN and Sorafenib and aged for 28 weeks. Mouse numbers for **b-d**, *Notch1^+/+^*, n=11; *Notch1*^+/-^ n=10; *Notch1*^-/-^, n=12. **b**. Esophagi from treated control, heterozygous, homozygous mice collected at experimental end point. Images are representative of each group. Scale bar, 1mm. **c**. Tumor density per genotype. Mean ± SEM, each dot represents a mouse, number of mice as shown in b. One-way ANOVA; adjusted p values from Tukey’s multiple comparisons test. **d.** Tumor areas per genotype. Mean± SEM, each dot represents a tumor. One-way ANOVA; adjusted p values from Tukey’s multiple comparisons test. **e, f**. Tumors from *Notch1^+/+^* and *Notch1^-/-^* mice were stained for immunostained for KRT14 (green) (**e**), or E-cadherin (CDH1, white) (**f**). Right hand images in **f** are magnified views of areas shown by a white square. White arrows indicate areas with reduced CDH1 staining. Images are representative of n=8 stained tumors from 6 mice of each genotype. Scale bars, 100 μm (**e, f** left panel) and 30 μm (**f** right panel). **g,h.** Mean E-cadherin intensity relative to DNA staining was quantified inside the tumor mass and compared to the corresponding adjacent normal tissue in wild type (**g**) and *Notch1^-/-^* (**h**) tissues (n=8 tumors from 6 mice for each genotype). Two tailed paired Student’s t-test. **i.** *YFPCreNotch1^+/+^* mice were treated with DEN/SOR and aged for 9 weeks. Mice were treated with anti-NRR1.1E3 or with control anti-Ragweed antibody (CTRL) for 6 weeks before collection. **j**. Representative tumors marked by KRT6a staining (red) are shown with white arrows in esophageal epithelium from control and anti-NRR1.1E3 treated mice. Scale bars: 100 μm. **k.** Quantification of the tumor area (Mean ± SEM, each dot represents a tumor, n=4 mice per group). P values from two-tailed unpaired Student’s t-test. AU, arbitrary unit.

These observations argue that *Notch1* promotes tumor growth. To further test this hypothesis, we treated wild type mice with a neutralizing antibody targeting the negative regulatory region (anti- NRR1.1E3) of NOTCH1, blocking its cleavage and NICD1 release ^34^. A short-term analysis revealed that the antibody was able to reduce levels of cleaved NOTCH1 in esophageal epithelium, abolish nuclear NICD1 immunostaining and normalize altered levels of multiple transcripts dysregulated by loss of *Notch1* such as *Sox9, Igfbp3, Gli2, Tgm2* and *Adam8 in vivo* **(Extended data Fig. 8a-e**). To determine the dosage of anti-NRR1.1E3 that would both inhibit NOTCH1 in esophageal epithelium and was well tolerated over an extended period of treatment we performed a clonal competition assay. *Notch1^-/-^* clones were induced in *YFPCReNotch1^flox/flox^* mice a week before a three-week course of antibody treatment (**Extended data Fig. 8f**). Effective blockade NOTCH1 signaling in the epithelium eliminates the competitive advantage of *Notch1^-/-^* clones over wild type cells, so mutant and wild type cells compete neutrally and mutant clone size will be reduced compared with controls (**Extended data Fig. 8g**). 10mg/kg Anti-NRR1.1E3 was both tolerated and significantly reduced in *Notch1^-/-^* clone size (p<0.0001, One-way ANOVA; Tukey’s multiple comparisons test) (**Extended data Fig. 8h,i**).

Wild type mice were given DEN and Sorafenib after which tumors were allowed to develop for 9 weeks. Animals were then treated with anti-NRR1.1E3 or control antibody at 10mg/kg for 6 weeks (**Fig. 7i**). The anti-NOTCH1 antibody treatment resulted in significant reduction in tumor size compared with controls, indicating active Notch signaling promotes the growth of established lesions (**Fig.7 j-k**).

## Discussion

These results shed light on the disparity in the prevalence of *NOTCH1* mutations in normal esophageal epithelium and tumors. A consequence of normal progenitor behavior is that most clones carrying mutations that do not alter progenitor cell behavior are lost by differentiation, making biallelic loss unlikely. Mutations reducing the function of one *Notch1* allele confer a competitive advantage on mutant progenitors, giving them a greatly enhanced probability of founding persistent, expanding clones. As the heterozygous mutant population grows, the likelihood that the remaining allele will be lost increases. When this happens, it confers a further increase in fitness. Over time, the result is extensive colonization of the epithelium by biallelic mutants, as wild type and heterozygous cells are displaced.

Once an area has been colonized by biallelic Notch1 mutants, the behavior of mutant progenitors reverts towards that of wild type progenitors so that the epithelium remains histologically normal in both humans and mice. Loss of *Notch1* does impact transcription of genes such as MCM proteins implicated in DNA replication and transcripts that regulate squamous epithelial morphogenesis, but functional and histological analyses indicate the epithelium is not substantially perturbed by these changes. These observations explain the remarkably high prevalence of *Notch1* mutations in normal appearing esophagus.

In contrast to normal epithelium, the loss of *Notch1* has a substantial phenotypic effect in carcinogenesis. *Notch1* is less mutated in tumors than wild type epithelium and its loss impairs tumor growth, an effect recapitulated by NOTCH1 neutralizing antibodies. How might wild type *Notch1* promote tumor growth? Our transcriptional analysis suggests some potential mechanisms. Examples include transcripts of genes that promote growth and invasiveness in ESCC cells, such as *Nrp2, Cdk14* and *Sox9* which are more highly expressed in *Notch1* wild type than null esophageal cells ^35–37^. Conversely, transcription of *Cldn7* which represses ESCC invasiveness is upregulated in *Notch1^-/-^* cells ^38,39^. Our study opens the way for further characterization of the pleotrophic effects of *Notch1* in esophageal tumors.

We conclude that *Notch1* illustrates how inactivating mutations in the same gene can drive clonal expansion in normal tissue but inhibit tumor growth. Our results further raise the possibility that blockade of NOTCH1 may have therapeutic potential in early esophageal neoplasia by reducing the growth of premalignant tumors. Multiple NOTCH1 inhibitors are in preclinical and clinical development and further investigation of their role in esophageal neoplasia seems warranted.

## Author contributions

E.A., J.C.F, and P.H.J designed experiments. E.A. and J.C.F performed experiments. E.A., S.C.D, M.W.J.H, S.H.O and R.S. analysed experimental and sequencing data. S.C.D. and M.G. created pipelines for copy number and scRNA-seq analyses. M.W.J.H. and B.A.H. performed clone simulations. C.W.S. provided neutralizing antibodies. E.A, S.C.D, M.W.J.H. and P.H.J wrote the paper. M.G, B.AH. and P.H.J supervised the research. All authors discussed the results and commented on the manuscript.

## Acknowledgements

This work was supported by grants from the Wellcome Trust to the Wellcome Sanger Institute (098051 and 296194) and Cancer Research UK Programme Grants to P.H.J. (C609/A17257 and C609/A27326). E.A. benefited from the award of a Postdoc fellowship, 2016-21, from the Wellcome Sanger Institute. B.A.H. and M.W.J.H. are supported by the Medical Research Council (Grant-in-Aid to the MRC Cancer unit grant no. MC_UU_12022/9 and NIRG to B.A.H. grant no. MR/S000216/1). M.W.J.H. acknowledges support from the Harrison Watson Fund at Clare College, Cambridge. B.A.H. acknowledges support from the Royal Society (grant no. UF130039). S.C.D. benefited from the award of an ESPOD fellowship, 2018-21, from the Wellcome Sanger Institute and the European Bioinformatics Institute EMBL-EBI.

We thank Esther Choolun, Tom Metcalf and Sanger RSF facility for technical support with animal research. We thank Yvette Hooks, Claire Hardy, Calli Latimer and staff from the CASM support laboratory for technical support with histology and sequencing.

## Legends for videos and tables

### Supplementary tables

The supplementary tables include source data for all figures

Supplementary Table 1: List of genes in the targeted sequencing experiments performed in this study

Supplementary Table 2a: Targeted sequencing of Human esophageal epithelium micro-biopsies; Complete mutation calling with Shearwater (part1) and from Caveman and Pindel pipelines (Part2)

Supplementary Table 2b: dNdS analysis of the Human micro-biopsy sequencing

Supplementary Table 2c: NOTCH1 mutations summary in Human esophageal epithelium micro- biopsies

Supplementary table 3: Human samples metadata

Supplementary table 4a: Notch1 mutations and CNLOH analysis in NOTCH1 stained *Notch1+/-* microbiopsies

Supplementary table 4b: Mutation calling in *Notch1+/-* micro-biopsies

Supplementary Table 5a: Differentially expressed genes in Notch1 mutant mouse esophageal epithelium

Supplementary Table 5b: EnrichGO analysis in *Notch1-/-* vs *Notch1+/+* and *Notch1+/-* vs *Notch1+/+* esophageal epithelium

Supplementary Table 5c: Differential expression analysis of canonical Notch targets and keratinocyte function related genes in *Notch1-/-* mouse tissue

Supplementary table 6: Single-cell transcriptomic analysis of *Notch1-/-* and *Notch1+/+* mouse esophageal epithelium

Supplementary table 7a: Mutation calling from targeted sequencing of DEN SOR treated esophageal epithelium

Supplementary table 7b: Mutation calling from targeted sequencing of esophageal tumors 28 weeks post-DEN and Sorafenib treatment

Supplementary Table 7c: dNdS analysis in normal mouse esophageal epithelium after DEN SOR treatment

Supplementary Table 7d: dNdS analysis in mouse esophageal tumors

Supplementary table 8: Antibodies used for immunostaining

#### Video1

In silico simulations showing clonal expansion in YFPCreNotch1 mutant mouse model over a period of 30 days after clonal induction. The top video (+/+) shows YFP+ *Notch1^+/+^* cells (green) expanding within a wild type epithelium (magenta). The middle video (+/-) shows YFP+ *Notch1^+/+^* and YFP+ *Notch1^+/-^* clones (green) as well as YFP- *Notch1^+/-^* (purple) evolving in a wild type tissue (magenta). The bottom video (-/-) shows YFP+ *Notch1^+/+^* and YFP+ *Notch1^-/-^* clones (green) as well as YFP-*Notch1^-/-^* (blue) evolving in a wild type tissue (magenta). 2-dimensional Wright-Fisher style model of clone dynamics in the esophagus and parameter fitting are described in **Extended data Fig.4** and **Supplementary Note**.

#### Video2

In silico simulations showing the effect of *Notch1* haploinsufficiency on the appearance of spontaneous *Notch1^-/-^* clone in the epithelium. The top video shows the scenario of *Notch1* haploinsufficiency (HI), using the best fit fitness parameter to the experimental analysis of *Notch1^+/-^*clones. The bottom video shows the scenario of *Notch1^+/-^* fitness being neutral (haplosufficiency, HS). *Notch1^+/-^* clones are shown in purple, *Notch1^-/-^* emerging by selection from *Notch1^+/-^* cells are shown in red and wild type cells are shown in light blue. Simulation results are shown over a period of 5000 days after the scattered induction *Notch1^+/-^* clones. In both cases, the best fit fitness parameter for *Notch1^-/-^* fitness was used. See **Extended data Fig.4** and **Supplementary Note**.

## Methods

### Human samples collection and processing

#### Collection

Esophageal tissue was obtained from deceased organ donors from whom organs were being retrieved for transplantation. Informed consent for the use of tissue was obtained from the donor’s family (REC reference: 15/EE/0152 NRES Committee East of England - Cambridge South). A full thickness segment of mid-esophagus was excised within 60 minutes of circulatory arrest and preserved in University of Wisconsin (UW) organ preservation solution (Belzer UW® Cold Storage Solution, Bridge to Life, USA) until processing. Tissue was flash frozen in tissue freezing medium (Leica catalog no. 14020108926).

#### Sampling for sequencin

For sequencing 20 μm thick cryosections were fixed with 4% paraformaldehyde for 10 min. Sections were stained for NOTCH1 (Cell Signaling catalog no. 3608) using the Rabbit specific HRP/DAB (ABC) Detection IHC Kit (Abcam Plc catalog no. ab64261) and counterstained with Haematoxylin. 250 to 1000 μm lengths of epithelium sections were microdissected from 4 to 6 successive serial sections and DNA extracted using a QIAMP DNA microkit (Qiagen) by digesting overnight and following manufacturer’s instructions. Flash frozen esophageal muscle DNA was used as the germline control and DNA extracted as for the epithelial samples.

#### Targeted sequencing

Samples were multiplexed and sequenced on an Illumina HiSeq 2000 sequencer using paired-end 75-base pair (bp) reads. Agilent SureSelect custom bait capture kits were used (**Supplementary table 1**). Details on DNA inputs, sequencing alignment, coverage metrics and pipelines used for mutation calling and copy number analysis are given in the **Supplementary Note.** Total number of protein altering mutations among the 60 sequenced samples mentioned in the main text are from CaVEMan and cgpPindel variant calling algorithms ^40,41^. *NOTCH1* mutations shown in **Fig.1c** were generated from these two algorithms with the addition of calls from ShearwaterML algorithm ^1,5,42^ which detects mutations with low variant allele frequency, see **Supplementary table 2c**.

#### Immunostaining and analysis of contiguous segments

For *in situ* quantifications, 10 μm thick serial cryosections were fixed with 4% paraformaldehyde for 10 min. For NOTCH1 and NICD1 detection and quantification **Fig. 1d-e**), three successive sections were stained respectively for NOTCH1/KRT4/ITGA6/DNA (1), NICD1/KRT14/DNA (2) and NOTCH1/KRT14/ITGA6/DNA (3) and imaged in their entirety. Corresponding areas in each section were identified based on tissue structure, distances and staining. Contiguous areas were analyzed for presence or absence of NOTCH1 or nuclear NICD1 staining and their length along the section measured using Volocity 6 software (Perkin Elmer). For morphological analysis of NICD1 and positive negative areas (**Extended data Fig.1 f-h**), sections were stained for NICD1/KRT14/DNA or NICD1/Ki67/DNA and epithelial thickness measured and cell counting performed using Volocity 6 software (Perkin Elmer). Cell density was measured by counting the number of nuclei identified across the width of the epithelium on high resolution images of cryosections stained with DAPI and keratinocyte markers such as KRT4 and ITGA6 and dividing this number by the measured surface area.

### Animals

All experiments were conducted according to the UK Home Office Project Licenses P14FED054 and PF4639B40. Male and female adult mice were used for in vivo experiments. Animals were housed in individually ventilated cages and fed on standard chow and maintained at SPOF health status. B6.129X1-*Notch1^tm2Rko/GridJ^* mice were purchased from the Jackson Laboratory and crossed with *Rosa26^floxedYFP^* and *AhCre^ERT^* to generate *YFPCreNotch1* triple mutant mice (**Extended data Fig. 2b-d**) ^13,17,43^. C57BL/6J wild-type mice were also used as indicated.

### qPCR Recombination assay

Genomic DNA from esophageal epithelium whole mount tissue was extracted using QIAamp DNA Micro Kit (Qiagen, catalog no. 56304). Specific primer sets were designed to analyze excision of the floxed exon1 of *Notch1* by *Cre* recombinase (**Extended data Fig. 3c**). Primer set A allows intragenic normalization using the non-floxed *Notch1* exon 3: Forward: GCCTGACACCTCTGGACAAC; Reverse: GCGGCACTTGTACTCTGTGA. Primer set B measures the disappearance floxed exon 1 with recombination: Forward: AGGAAAGAGGGCATCAGAGG; Reverse: GGACGTTGACGCACACCA. Quantitative PCR on genomic DNA was carried out using specific primers and SYBR Green master mix (Thermo Fisher Scientific, catalog no. 4309155) according to the manufacturer’s instructions in a StepOnePlus™ Real-Time PCR System (Thermo Fisher Scientific, catalog no. 4376600). Relative q-PCR expression was calculated using delta-delta Ct method, a wild type reference sample was used within the same assay. Validation of the linearity of the recombination assay was performed against a standard curve reproducing different recombination rates with Exon1/Exon3 ratio of 1, 0.75, 0.5, 0.25 and 0. The Standard curve was made using diluted genomic DNA from the esophagus of highly induced and fully recombined *Notch1^-/-^* mice (as verified by qPCR, staining and protein assay) and from *Notch1* wild type tissue. Full recombination of the esophageal epithelium will reduce the Exon1/Exon3 ratio to zero in induced *Notch1^-/-^* mice and to 0.5 in induced *Notch1^+/-^* mice (**Extended data Fig. 3e, Extended data Fig.6a**).

### RT-qPCR assay

RNA extractions were performed on mouse esophageal tissue as described in the RNA sequencing method section. Total RNA was measured using Qubit™ RNA BR Assay Kit (Thermo Fishcer Scientific, catalog no. Q10211). cDNA synthesis of 500ng total RNA was performed using QuantiTect Reverse Transcription Kit (Qiagen, catalog no. 205313). RT-qPCR was performed with Taqman Fast Advanced Master Mix (Thermo Fisher Scientific, catalog no. 4444557) on StepOnePlus™ Real-Time PCR System (Thermo Fisher Scientific, catalog no. 4376600) and analyzed using StepOne Software v2.3. Relative q-PCR expression to *Gapdh* housekeeping gene was calculated using delta-delta Ct method. The specific Taqman primers u*sed* for quantification were the following*: Gapdh: Mm99999915_g1; Notch1: Mm00627185_m1; Gli*2: Mm01293117_m1; *Igfbp3*: Mm01187817_m1; *Adam8*: Mm00545745_m1; *Tgm2*: Mm00436979_m1; *Sox9*: Mm00448840_m1.

### Immune Capillary Electrophoresis

RLT Plus lysates with Complete Protease Inhibitor (Roche, catalog no. 11836170001) homogenized as described in the ‘RNA sequencing’ section were passed through the RNA binding column from the AllPrep DNA/RNA Mini kit (Qiagen) and the flow through was collected for protein precipitation. For precipitation, 9 volumes of ice cold pure Ethanol were mixed to the lysates before storage overnight at -80 degrees. Precipitates were spun 30min at 20,000 g at 4 degrees, pellets were dried and solubilized progressively with 5% Sodium dodecyl sulfate in 100mM TEAB solution (Sigma-Aldrich, catalog no. T7408). Total protein quantification was performed using Pierce™ BCA Protein Assay Kit (Thermo Fisher Scientific, catalog no. 10678484)). Immune capillary electrophoresis was performed using Wes Simple^TM^ (ProteinSimple) following manufacturer’s instructions and analyzed using Compass for SW version 4.1.0. Primary antibodies were the following: anti-NOTCH1 targeting C terminus of the protein (Cell signaling, catalog no. 3608); anti-NOTCH2 targeting C terminus of the protein (Cell signaling, catalog no. 5732); alpha-TUBULIN (Cell signaling, catalog no. 2125).

### Whole-mount sample preparation of mouse esophagus

Mouse esophagus was cut and opened longitudinally before removing the muscle layer with forceps. For the clonal lineage tracing experiments, the tissue was incubated for 15 minutes in Dispase I (Roche catalog no. 04942086001) diluted at 1mg/ml in PBS before separating the epithelium with fine forceps. For all the other immunostaining experiments, tissue was incubated for 2h15 to 3 h in 5 mM EDTA at 37 °C before gentle peeling of the epithelium. The epithelium was flattened and fixed in 4% paraformaldehyde for 1h15 at room temperature under agitation. Tissues were then washed in PBS and stored at 4 °C.

### Whole mount immunostaining

Whole mount tissues were blocked an hour in staining buffer (0.5% BSA, 0.25% fish skin gelatin, 0.5% Triton X-100, 10% donkey serum in PHEM). Samples were incubated with primary antibodies (**Supplementary Table 8**) in staining buffer overnight at room temperature, followed by three washes of 30 min with 0.2% Tween-20 in PHEM. Samples were incubated with secondary antibodies (**Supplementary Table 8)** in staining buffer for 3h at room temperature and washed. Finally, tissues were incubated overnight at room temperature with 1 μg.ml^−1^ DAPI or 0.5 μM Sytox™ Blue solution (Biolegend, catalog no. 425305, see section on ‘preparation of mouse samples for sequencing’) to counterstain cell nuclei and mounted using Vectashield mounting media (Vector Laboratories, catalog no. H-1000).

#### Cryosections

Esophageal tissue was flash frozen in tissue freezing medium (Leica catalog no. 14020108926). 10 µm transversal sections were fixed with 4% paraformaldehyde for 10 min, blocked in staining buffer (0.5% BSA, 0.25% fish skin gelatin, 0.5% Triton X-100, 10% donkey serum in PHEM) and stained with the respective primary and secondary antibodies for 3 hours to overnight at room temperature (**Supplementary Table 8).** Samples were washed with PHEM buffer between incubations. Prior to NICD1 staining, sections were incubated 20 minutes in 50mM Glycine/PBS solution. Finally, tissues were incubated overnight at room temperature with 1 μg.ml^−1^ DAPI or 0.5 μM Sytox™ Blue solution (Biolegend, catalog no. 425305) to counterstain cell nuclei and mounted using Vectashield mounting media (Vector Laboratories, catalog no. H-1000).

### Histology

Hematoxylin and eosin staining was either performed on 10 µm cryosectioned tissue processed as described above or on 5 µm paraffin embedded tissue sections. Prior paraffin embedding, esophageal tissue was collected and fixed in 4% paraformaldehyde for at least 2 hours before undergoing progressive dehydration in Tissue-Tek VIP® 6 AI tissue processor (Sakura). Slides were then scanned at objective 20x using NanoZoomer S60 Digital slide scanner (Hamamatsu).

### Confocal microscopy

Immunofluorescence images were acquired on a Leica TCS SP8 confocal microscope using ×10, ×20 or ×40 objectives. Typical settings for acquisition were optimal pinhole, line average 3-4, and scan speed 400-600 Hz and a resolution of 1024 × 1024 pixels. Visualization and image analysis were performed using Volocity 6 Image Analysis software (PerkinElmer).

### Lineage tracing using a YFP reporter

To genetically label wild type and *Notch1* mutant clones in the same tissue, we crossed floxed *Notch1* mice with floxed *Rosa26^EYFP^* animals carrying a conditional fluorescent YFP reporter at the *Rosa26* locus and the inducible *AhCre^ERT^* mice ^13,17^. Pharmaceutical treatment with β-naphthoflavone (BNF, MP Biomedicals, catalog no.156738) and tamoxifen (TAM, SigmaAldrich, catalog no. N3633) allows the expression and nuclear translocation of the Cre protein and the subsequent recombination of the floxed *Notch1* resulting in the excision of its first exon and/or *Rosa26* alleles resulting in the expression of the fluorescent reporter YFP. Titration of the dosage of BNF/TAM was performed to allow recombination of in a small proportion of the esophageal basal cells, called clonal level. This allows the recombined cells to expand over time and generate clones without fusion. Excision of the *Notch1* allele and expression of the YFP reporter at the *Rosa26* locus can occur in the cells either in combination or separately, resulting in four different population of cells, N*otch1* mutant or not and expressing YFP or not (**Extended data Fig. 2b, c**). Mice between 10-16 weeks of age were injected intraperitoneally (i.p.) with BNF (80 mg.kg^-1^) and TAM (0.125 mg) and collected at different time points. Staining and processing of the whole mount tissue allowed us to track the clones based on YFP and NOTCH1 expression (**Extended data Fig. 3**). YFP expressing clones were imaged with confocal microscopy at 40x objective. In induced *Notch1^flox/flox^* tissue, YFP+ clones expressing NOTCH1 were categorized as wild type, while the clones without detectable expression of NOTCH1 were classified as *Notch1^-/-^*.

In order to distinguish the wild type and *Notch1+/-* clonal populations in induced *Notch1^+/floxed^* tissue, we set up a method based on measurement of NOTCH1 staining. For each clone, the mean intensity of NOTCH1 immunofluorescence was recorded in the basal cells expressing the YFP reporter, normalized to the mean intensity of the corresponding DNA stain in each image. We analyzed the NOTCH1 staining intensity distribution in 3 wild type mice in n=126 YFP+ clones and used this as a reference (**Extended data Fig. 3h**). The distribution of the relative intensities could be approximated with a normal curve (p=0. 25, Shapiro-Wilks test) with mean 1.02 and standard deviation 0.15, fit with Python package Scipy v1.5.2. The distribution of relative NOTCH1 staining intensity of the n=172 YFP clones in induced *Notch1^+/flox^* mice was significantly different from the distribution in the *Notch1^+/+^* mice (p=5e^-27^, two-tailed Kolmogorov-Smirnov test). To confirm that the clones in induced *Notch1^+/flox^* mice were a mixture of *Notch1* wild type and *Notch1^+/-^* clones, we fit a Gaussian Mixture model with two components to the *Notch1^+/flox^* relative intensity distribution using the Python package Scikit-learn v0.23.2. The resultant fit consisted of a normal distribution highly similar to that fit to the clones from the *Notch1^+/+^* mice (mean=1.0, sd=0.15), and a normal distribution with half the intensity of the *Notch1* wild type clones (mean=0.53, sd=0.15). We chose to split the induced *Notch1^+/-^* clones using a relative intensity threshold of 0.75. This threshold is slightly lower than the mid-point of the two distributions, and was selected so that very few of the wild type clones (<5%) would be categorized as *Notch1^+/-^.* The higher intensity clones were assumed to be *Notch1* wild type (+/+ in +/flox) and had a similar distribution of staining intensity to those in the *Notch1^+/+^* mice (p=0.38, two-tailed Kolmogorov-Smirnov test). The remaining clones were categorized as *Notch1^+/-^* (**Extended data Fig. 3e**). These methods were used at each time point. Number of basal and suprabasal cells in each clone was counted using Volocity 6 Image Analysis software (PerkinElmer).

Clones with no basal cells in the induced mutant mice could not be categorized with this method but represented only ∼1% of the clones in the dataset. Fusion of expanding clones prevented quantification of clone size beyond 4 weeks in induced Notch1^flox/flox^ esophagus and beyond 13 weeks in induced Notch1^+/flox^ tissue.

### EDU lineage tracing

EdU (5-ethynyl-2ʹ-deoxyuridine) incorporates during replication in the proliferating cells located in the basal layer of the esophageal epithelium (**Extended data Fig. 2a**). EdU i.p. injection at 10 µg was performed either at 1 hour or 48h before tissue collection. Tissue was processed and EdU was detected in whole mount using Click-iT EdU imaging kit (Life Technologies, catalog no. C10338 or C10340). Images were acquired by confocal microscopy and analyzed using Volocity 6 software.

Performed 1 hour before collection, the assay was used to quantify the proliferation rate in the esophageal epithelium. The proportion of EdU+ basal cells: total basal cells was recorded as a measure of proliferation. Performed 48h before collection, this assay was used to quantify differentiation. Indeed, the EdU labelled cells generate daughter cells that can either stay in the basal layer or differentiate and leave the basal layer. The proportion of EdU positive suprabasal cells relative to total EdU+ cells was recorded as a measure of cell stratification. In the clonal induction experiment, the wild type neighbor cells of the mutant clones called +/+ edge were identified as the first row of cells adjacent to the YFP expressing cells while the distant cells called +/+ distant were randomly situated further away from the clone (**Fig.2 g-n**).

### Aging experiments

*YFPCreNotch1* mice between 10-16 weeks of age were injected intraperitoneally (ip) with BNF at 80 mg.kg^-1^ and TAM at 1 mg for *Notch1^+/+^* and *Notch1^+/flox-^* and at 0.25mg for *Notch1^flox/flox^* and collected at different time points after induction and up to 78 week-old. (**Fig. 3c, Fig.5, Extended data Fig.5**). *Notch1^+/+^* or non-induced mice were used as wild type controls as indicated. A lower dose of Tamoxifen was used for the *Notch1^flox/flox^* mice in order to minimize the recombination of the *Notch1* allele in the corneal epithelium, possibly leading to corneal opacification and keratinization ^44^.

### Projected NOTCH1 stained area quantification

To quantify the percentage of NOTCH1 positive or negative area in the entire esophageal epithelium or the projected surface of NOTCH1 negative clones (**Fig. 3b, Fig. 6e, Extended data Fig. 8i**), whole mount esophageal epithelia were prepared and stained for NOTCH1 and counterstained with DAPI or Sytox™ Blue as described in the dedicated sections. Tissues were imaged entirely using high precision motorized stage coupled to a Leica TCS SP8 confocal microscope to obtain contiguous 3D images of all epithelial layers and merged using the mosaic function of the Leica software. Typical settings for acquisition of multiple z stacks were optimized 2.41 μm z step size, zoom ×1, optimal pinhole, line average 4, scan speed 400-600 Hz and a resolution of 1024 × 1024 pixels using an ×10 HC PL Apo CS Dry objective with a 0.4 numerical aperture. Images were processed using Volocity 6 software. To measure their projected surface area, images were opened using the ‘extended focus’ visualization mode on the Volocity software. Regions of interest were defined with ROI tool allowing surface area measurement. NOTCH1 staining was automatically detected based on defined intensity and minimum object size.

### Detection of spontaneous mutant clones and *ex vivo Notch1* mutant functional assay in aging *Notch1^+/-^* tissue

Esophageal tissue from induced *YFPCreNotch1^+/-^* mice aged to 54-78 week-old was incubated for 2h30 in 5mM EDTA solution at 37 °C before peeling. Whole mount esophageal epithelium was stained for NOTCH1/ YFP and Sytox™ Blue and imaged. In the aged *YFPCreNotch1^+/-^* tissues, the detection of large and ovoid areas negative for NOTCH1 staining allowed the identification of putative spontaneous NOTCH1 negative clones (**Fig. 3b, e**). To identify putative clones expressing NOTCH1, we used the combination of NOTCH1 and YFP staining. In highly induced heterozygous tissue, YFP staining normally appear as a patchwork of small YFP+ clones. Putative clones appeared as large and ovoid areas devoid or fully stained with YFP (**Fig.3e**). Sequencing confirmed that this method allowed the identification of large spontaneous clones carrying *Notch1* mutations on the non-recombined allele (**Fig. 3e-g**). EDTA treatment triggers the cleavage of NICD1, followed by nuclear translocation without ligand activation^20^. We used this assay to test the functionality of the cleavage of the spontaneous NOTCH1 positive clones identified (**Extended data Fig. 5e-j**).

### Carcinogen treatment

Mice were induced with BNF/TAM at a dose that allowed full coverage of the tissue with the mutant *Notch1* heterozygous or homozygous cells within three months. *YFP*C*reNotch1^+/flox^* mice were injected intraperitoneally (i.p.) on two consecutive days with BNF at 80 mg.kg^-1^ and TAM at 1 mg for and YFPCre*Notch1^flox/flox^* were injected once with BNF at 80 mg.kg^-1^ and TAM at 0.25 mg. Non-induced *YFPCreNotch1* mice were used as controls. Then, in order the generate mutations and tumor formation in the esophageal epithelium, mice were treated with DEN (Sigma, catalog no. N0756) in sweetened drinking water (40 mg/L) for 24 h, 3 days a week (Monday, Wednesday and Friday) for 8 weeks ^16,19^. Sorafenib (LC Chemicals, catalog no. S8502) was then administered at 50mg/kg (5 microliters of 10mg/ml solution per g bodyweight) by i.p. injection on alternate days during 6 weeks, for a total of 21 doses ^29^.

Mice were aged for 28 weeks after the last dose of Sorafenib and esophageal tissue was collected and opened longitudinally. Macroscopic images of unpeeled tissue were obtained under Leica M80 zoom stereomicroscope with Leica Plan 1.0x Objective M-Series 10450167 coupled with Leica DFC295 Camera (Leica microsystems). Macroscopic tumors were punched and flash frozen in tissue freezing medium (Leica catalog no. 14020108926) for further processing. Esophageal tissue was peeled for 3 hours in 5mM EDTA, fixed, processed through whole mount immunostaining for the Keratin 6a stress marker and DNA staining before confocal imaging. Detailed analysis of macroscopic and microscopic images was performed to quantify the projected areas of the lumps using ROI tool in Volocity 6 software.

### Antibody treatment validation

Antibody validation was performed in two steps. For step 1, C57BL/6J were injected intraperitoneally at high dose with rat anti-Notch1 NRR hybridoma Clone 1E3.19.1 (anti-NRR1.1E3, Genentech) at 25mg.kg^-1^ (n=3 mice per group). Anti-NRR1.1E3 is distinct from the anti-NRR1 from Wu et al. ^34^. This rat monoclonal antibody has a lower affinity for mouse anti-NRR and is better tolerated. Antibody Ragweed:9652 10D9.W.STABLE mIgG2a (CTRL, Genentech) was used as control at 25mg/kg^-1^. 3 days later, esophageal tissue was collected and processed for RT-qPCR assay and protein quantification as described in the ‘RT-qPCR’ and ‘Immune Capillary Electrophoresis assay’ (ICE) sections. At the protein level, cleaved transmembrane/intracellular regions (NTM + NICD) of NOTCH1 and NOTCH2 were quantified by ICE. The assay confirmed the specificity of the antibodies for NOTCH1 neutralization (**Extended data Fig.8a-c**). Tissue was also sectioned and stained for NICD1, confirming the absence of active NOTCH1 in basal keratinocytes with the anti-NRR1.1E3 treatment **(Extended data Fig.8d**). At the transcriptional level, RT-qPCR for *Notch1* loss of functions markers Igfbp3, Tgm2, Gli2, Adam8 and Sox9 were performed (**Extended data Fig.8e**). For step 2, treatment doses were titrated to allow efficiency and safety of the treatment over an extended period of time. *YFPCreNotch1^flox/flox^* mice were induced at clonal level, and starting a week later, were treated once a week with the antibodies for 3 weeks and tissue was collected, peeled and whole mount stained for NOTCH1. As expansion of clones is subject to the fitness of the neighbor cells ^19^, a total NOTCH1 blockade would be expected to neutralize the competitive advantage of *Notch1^-/-^* clones over wild type cells, reducing clonal expansion. NOTCH1 negative staining areas were therefore measured as to determine the efficacy of the antibody in inhibiting NOTCH1 in esophageal epithelium (**Extended data Fig. 8f-i**). Anti-NRR1.1E3 at 10mg.kg-1 offered the strongest reduction in NOTCH1 negative clone size showing the efficacy of the treatment. The health and weight of the mice was assessed through the treatment. n=4 mice injected with Ragweed controls at 10mg/kg; n=3 mice treatedNRR1.1E3 at 10mg/kg; n=2 mice given NRR1.1E3 10mg/kg loading dose (week 1: 10mg/kg, week 2: 7.5, week 3: 5 mg/kg); n=1 mouse given NRR1.1E3 5mg/kg; and n=1 mouse given NRR1.1E3 5mg/kg loading (week1: 5mg/kg, week 2: 4mg/kg, week 3: 3mg/kg). Antibody NRR1.1E3 did not lead to any weight loss or observable adverse effects.

### Immunotherapeutic treatment

*YFPCreNotch1^+/+^* mice were first treated with DEN and Sorafenib and aged for 9 weeks to allow the tumors to start growing before starting a treatment with anti-NRR1.1E3 at 10mg.kg-1 or with Ragweed control at 10mg.kg-1 (n=4 mice per group), once a week for 6 weeks (**Fig. 7i-k**). One week after the last dosage, tissue was collected and processed for macroscopic and microscopic quantification of the projected areas of the tumors using Volocity 6 software as described in ‘carcinogen treatment’ section

### Cell density in mouse esophagus

For aged mouse tissue (**Fig. 5c**), density of the basal cells was measured on whole mount stained tissue, imaged at 40x objective using Leica TCD SP8 confocal microscope (see ‘confocal microscopy’). DAPI+ or Sytox™ Blue + basal nuclei were quantified per area in 3 to 6 random positions of the tissue, n=4 mice per genotype analyzed.

### Cell counting and epithelial thickness in mouse tissue cryosections

Esophageal epithelial thickness was quantified in transversal cryosections stained with Hematoxylin and eosin (H&E) with NanoZoomer Digital Pathology software (NDP.view2, Hamamatsu). Measurements were performed at 18 to 23 positions in the tissue and averaged for each mouse (n=3). For cell counting in cryosections, stained sections were imaged using Leica TCD SP8 confocal microscope at 40x objective as described in ‘confocal microscopy’ and analyzed with Volocity 6 software (Perkin Elmer). For counting Ki67 positive basal cells in the aged tissue (**Fig.5d**), tissue was stained for Ki67 & DNA and analysis were performed at three different positions per animal and averaged for each mouse (n=3-4). For Ki67 counting in the tumors (**Extended data Fig.7f**), tissue cryosections were stained for Ki67, KRT14 and DNA to identify keratinocytes in the tumor mass (N=8- 9 tumors from n=5 Notch1^+/+^ and 6 Notch1^-/-^ mice respectively). For NICD1 counting, sections of tumors and adjacent normal tissue were stained for NICD1, KRT14 and DNA. Proportion of KRT14+ keratinocytes expressing nuclear NICD1 inside the tumor mass and in the adjacent normal epithelium was quantified (n=10 tumors from 6 mice).

### Projected E-cadherin intensity quantification

Esophageal sections carrying tumors and adjacent normal tissue were stained for E-cadherin, Keratin 14, counterstained with DAPI **(Fig. 7f-h**) and imaged at high resolution using Leica TCD SP8 confocal microscope as described in the previous sections. To measure the intensity of the staining on projected areas, images were opened using the ‘extended focus’ visualization mode on the Volocity 6 software. Regions of interest (adjacent normal epithelium and keratinocytes populated tumor mass) were defined with ROI tool and mean intensity within areas were quantified. Mean intensity of E-cadherin was normalized on the mean intensity of DAPI for each ROI. Analysis compared measures in the tumors to the measures in the adjacent normal tissue (8 tumors from 5 wild type mice; 8 tumors from 6 *Notch1^-/-^* mice). We verified that E-cadherin staining was not affected in the normal tissue of DEN/SOR treated *Notch1^-/-^* mice compared to wild type mice using this method on tissues embedded in the same OCT bloc and stained together on the same slide (n=3 mice).

### RNA sequencing

*YFPCreNotch1^flox/flox^* and *YFPCreNotch1^+/flox^* mice between 10-16 weeks of age were injected intraperitoneally (i.p.) on two consecutive days with BNF at 80 mg.kg-1 and TAM at 1 mg and aged for 8 weeks before collection (**Fig. 4a).** The dose was set in allow *Notch1^+/-^* or by *Notch1^-/-^* cells to fully occupy the tissue by two months post induction, as measured by qPCR recombination assay (**Extended data Fig.3e**), qPCR assay and protein assay (**Fig.2b,c**). Uninduced littermates were used as wild type control. Esophageal tissues were incubated for 15 minutes in Dispase I (Roche catalog no. 04942086001) diluted at 1mg/ml in PBS before separating the epithelium with fine forceps. Peeled epithelium were lysed in RLT Plus buffer using MagNA Lyser Green Beads (Roche, catalog no. 03358941001) and FastPrep 24 homogenizer (MP Biomedicals, catalog no.116004500). Lysates were then processed for RNA and DNA extraction using AllPrep DNA/RNA Mini kit (Qiagen), with Complete Protease Inhibitor (Roche, catalog no. 11836170001) added instead of the proteinase K step, allowing protein extraction as described in the Immune Capillary Electrophoresis section. Total RNA was measured using Qubit™ RNA BR Assay Kit (Thermo Fisher Scientific, catalog no. Q10211).

For RNA-seq, libraries were prepared in an automated fashion using an Agilent Bravo robot with aKAPA Standard mRNA-Seq Kit (KAPA BIOSYSTEMS). In-house sequencing adaptors were ligated to 100-300 bp fragments of input dsDNA. Samples were then subjected to 10 PCR cycles using sanger_168 tag set of primers and paired-end sequencing was performed on Illumina HiSeq 2500 with 75 bp read length. Reads were mapped using STAR 2.5.3a, the alignment files were sorted and duplicate-marked using Biobambam2 2.0.54, and the read summarization performed by the htseq-count script from version 0.6.1p1 of the HTSeq framework ^45,46^. Differential gene expression was analyzed using the DESeq2 R package ^47^ and the downstream pathway analysis and visualization using R (https://www.R-project.org/) and the packages Pheatmap (https://cran.r-project.org/package=pheatmap), RColorBrewer (https://cran.r-project.org/package=RColorBrewer), clusterProfiler ^48^ and org.Mm.eg.db (https://bioconductor.org/packages/org.Mm.eg.db/). Background noise was filtered out from the dataset using the following rule: genes with an eigenvector value of >0.05 or <-0.05 for both PC1 and PC2 among wild type control samples were removed from the dataset (151 genes out of the initial 47,467 genes) for final analysis. Differentially-expressed genes (DEG) are hits reported by DESeq2 with adjusted p value (p_adj) of less than 0.05.

Heatmaps were generated from the ratio of TPM values of the treated sample over the average of the respective control samples. The volcano plots are plotted using the R package plotrix to enable gapped axis (https://cran.r-project.org/package=plotrix) using the output from DESeq2 as input.

### Single cell RNA sequencing

*YFPCreNotch1^flox/flox^* mice were injected intraperitoneally (ip) with BNF at 80 mg.kg^-1^ and TAM at 1 mg and aged for 11 weeks. Uninduced littermates were used as wild type controls. Mouse esophagus was collected and placed immediately in PBS on ice. Esophagus was opened and cut into 4 equal pieces and placed in 1ml dispase (2.4U/ml) at 37C for 10mins. Epithelia was then dissected away from the underlying stroma and muscle. Resulting epithelia was minced very finely using a scalpel and the slurry placed in 1ml trypsin-EDTA for 30mins at 37C. After incubation each sample was placed in 9ml ice-cold 0.04% BSA/PBS in a C-type GentleMacs tube (Miltenyi Biotec). Tissue was further dissociated using a GentleMACs dissociator with programs A1 and two rounds of program B1 with the sample kept on ice between programs. Samples were briefly centrifuged and filtered through a 30um filter (Miltenyi Biotec). Samples were then centrifuged at 4°C for 10mins at 500xg. The pellet was resuspended in ice-cold 0.04% BSA/PBS and centrifuged again a 4°C for 5mins at 300xg. Pellet was resuspended in ice-cold 0.04% BSA/PBS. Cells were counted and sufficient volume was placed into a Chromium reaction to allow 5000 cell recovery. Single cells were collected using Chromium (10X) with version 3 chemistry following manufacturer’s instructions with one mouse esophagus per inlet. cDNA was sequenced using HiSeq4000 (Illumina) with the following read length 28 bp read 1; 91 bp read 2; 8 bp index 1; 0 bp index 2 with a single inlet per lane. Details of analytic methods for sc-RNAseq are given in the **Supplementary Note**.

### Mouse sample preparation for DNA sequencing

Prior to sampling of the normal esophageal tissue, esophageal epithelium was processed for whole mount immunostaining and imaged with confocal microscopy as described in the corresponding sections. Tissue for sequencing was incubated for 1 hour with 0.5 μM Sytox™ Blue solution (Biolegend, catalog no. 425305) to counterstain cell nuclei. For sequencing, DNA from the ears of the same mice was extracted with the same method and used as germline controls.

#### Punching

For subclonal analysis of stained areas of interest, stained and imaged tissue was flattened and punched under a Fluorescent Stereo Microscope Leica M165 FC (Leica) using 0.25 mm diameter punch (Stoelting, catalog no. 57391) (**Fig. 3e, Extended data Fig.5**). DNA from the microbiopsies was extracted using Arcturus® PicoPure® DNA extraction kit (Applied Biosystems, catalog no. 11815-00) following the manufacturer’s instructions).

#### Gridding

For analysis of the mutations in the healthy esophageal epithelium, the esophageal epithelium was flattened, mapped and cut into 2 mm^2^ contiguous biopsies (**Fig. 6a-c, Extended data Fig. 8d-f)**. Samples were digested and DNA extracted using the QIAMP DNA microkit (Qiagen, catalog no. 56304) following the manufacturer’s instructions.

#### Tumor sampling

*Flash frozen* tumors collected as described in the ‘carcinogenesis’ section were cut into serial 50 µm cryosections and stained for KRT14, KRT6a and Sytox™ Blue (**Fig. 6a-c, Extended data Fig. 7a,b)**. In order to separate the tumor mass from the surrounding normal tissue and the underlying submucosa, sections were micro-dissected under a Fluorescent Stereo Microscope Leica M165 FC (Leica) using a micro knife (FST, catalog no. 10317-14). Dissected tumor from serial sections were pooled and tumor DNA was extracted using the QIAMP DNA microkit (Qiagen, catalog no. 56304) following the manufacturer’s instructions.

#### Targeted Sequencing

Samples were multiplexed and sequenced on Illumina HiSeq 2000 sequencer using paired-end 75-base pair (bp) reads. Agilent SureSelect custom bait capture kits were used (**Supplementary table 1**). Details on the custom bait kit, DNA inputs, sequencing alignment and coverage metrics are given in the **Supplementary Note.**

Details of processing and analytic methods for mutation calling and copy number alteration analysis are given in the Supplementary Note.

### Modelling

Stochastic simulations of clonal dynamics are explained in **Supplementary Note**.

### Statistical analysis

Data are expressed as mean values ± SEM unless otherwise stated. Statistical tests are indicated in figure legends. No statistical method was used to predetermine sample size. Data are expressed as mean values ± SEM unless otherwise stated. Statistical tests are indicated in figure legends. No statistical method was used to predetermine sample size. Animals of the correct genotype were randomly assigned to experimental groups. The investigators were not blinded to allocation during experiments and outcome assessment.

### Data availability

Accession numbers for the datasets are as follow:-Targeted sequencing of Human esophageal epithelium microbiopsies (EGA): EGAD00001006969. -Targeted sequencing of aged Notch1 +/- mouse esophageal epithelium microbiopsies (ENA): ERP126992-Targeted sequencing of mouse normal esophageal epithelium 28 weeks after DEN SOR treatment (ENA): ERP126993 -Targeted sequencing of mouse esophageal tumours 28 weeks after DEN SOR treatment (ENA): ERP126994-Transcriptomic analysis of Notch1 mutant esophageal epithelium (ENA): ERP126995 -Single cell transcriptional analysis of Notch1 mutant esophageal epithelium (ENA): ERP126996.

### Code availability

The codes developed in this study has been made publicly available and can be found at https://github.com/PHJonesGroup/Abby_etal_SI_code

### Biological Resources

Mouse strains are available from the Jax repository (https://www.jax.org), with the exception of the *Ahcre^ERT^* line which may be obtained by contacting the corresponding author.

## Supplementary Note

This supplementary note details the computational modelling used in this study to simulate clonal dynamics followed by a description of the DNA and sc-RNA sequencing approaches and pipelines used.

### 1. Stochastic simulations of clonal dynamics

#### 1.1. Previous models of mutant clonal dynamics in esophageal epithelium

Murine esophageal epithelium (EE) consists of layers of keratinocyte cells and is maintained by the proliferation of cells in the basal layer (**Extended data Fig. 2a**). Each basal cell division can form a pair of cells that remain in the basal layer and go on to divide, a pair of cells that will differentiate and stratify into the suprabasal layers without dividing, or one cell of each fate. The outcome of each division is random, but in homeostasis the balance between dividing and differentiating cells is maintained (**Extended data Fig. 2a**) ^1^ so that the overall number of proliferative cells remains constant. Due the random (stochastic) nature of the process, some cell lineages will be lost from the tissue and others will form large clones of the original cell ^1,2^.

A genetic mutation in a basal cell may introduce a bias towards the production of more proliferating cells, leading to the non-neutral expansion of a mutant clone ^3,4^. However, in normal EE, a clone cannot not expand indefinitely ^5,6^. Mutant clones compete for the limited space in the tissue, and expanding clones revert towards more neutral growth once they encounter similarly fit clones ^6^. Therefore, the fate of a cell division depends not only on the dividing cell, but also on the neighboring cells ^6^. This spatial clonal competition in EE has been previously been modelled using stochastic simulations of cellular automata on fixed 2D grids ^6^.

#### 1.2. 2D Wright-Fisher simulations

In this study, we wished to compare the growth of wild type (+/+) clones, induced heterozygous *Notch1^+/-^* clones and induced homozygous *Notch1^-/-^* clones. We were looking for broad differences between growth rates, not for precise characterization of the clonal dynamics of each genotype. We therefore aimed to keep the model simple, with few free parameters, so that the results of parameter fitting for each genotype could easily be compared.

We used simulations based on a Wright-Fisher model ^7^ constrained to a 2D grid ^8^. Each cell, *C_a,b_*, in the grid represents a cell in the basal layer of the EE. The subscripts denote the cell in location *a* in the grid and generation *b* of the simulation. We used a fixed size (500×500 cells) hexagonal grid with periodic boundary conditions.

Each cell has a fitness, *F_a,b_*. In each step of the simulation, each cell *C_a,b_* in the new generation picks a parent cell from its immediate neighborhood 𝒩*_a_* (the six adjacent cells plus the cell in the same grid position) in the previous generation (**Extended data Fig. 4a**). The chance of a cell being picked as the parent depends on its fitness relative to the other potential parent cells in the neighborhood,

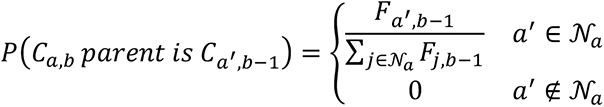

A cell can produce multiple offspring in the next generation and the offspring cells inherit the fitness of their parent cells. In this way, a clone with high fitness can expand over multiple generations (**Extended data Fig. 4a**). At the start of a simulation, a small proportion of cells in the grid were randomly selected to be the induced mutant cells. In each simulation, a single fitness value was assigned to all the mutant cells (see **section 1.4** below). The rest of the cells in the simulation were given a fitness of 1. We tracked the sizes of the clones that grew from the mutant cells.

Division rate did not significantly differ between the wild type and *Notch1* mutant tissue (**Fig. 2g-j**). We therefore fixed the division rate for all simulations at the division rate previously measured in the murine EE, 0.27 divisions per day ^1^. The fitting results showed that this was appropriate for capturing the average growth of the wild type clones in this model (see **section 1.4** below).

For each simulation, we had two parameters that were not fixed: the fitness of mutations and the induced proportion of cells at the start of the simulation. The induction proportion must be considered because as the clones expand they can collide and restrict each other’s growth. In a highly induced tissue, fit mutant clones will have less room to expand and will be smaller at later time points than in a sparsely induced tissue. The other parameter, mutant fitness, is the key parameter we wish to compare between the genotypes. It represents how much of a growth advantage the mutant has compared to the surrounding non-mutant cells.

#### 1.3. Approximate Bayesian Computation

To fit the parameters to the data we used Approximate Bayesian Computation (ABC) ^9,10^. In this fitting method, prior distributions for the parameters are defined. We used uniform distributions across wide intervals: mutant fitness between 0 and 50, induction between 0% and 10% of cells. A parameter set is randomly drawn from the prior distributions, and a simulation run with those parameters. The simulation results are measured against the observed data using summary statistics. We used the Kolmogorov-Smirnov statistic ^11^ to compare the simulated and experimentally observed basal clone size distributions. We used 100 simulated clones at each time point, and summed the Kolmogorov-Smirnov statistics calculated for each time point to get the total “distance” between the simulation and the data. If the distance between the simulation and the experimental data is less than a set threshold, those simulation parameters are accepted, otherwise they are rejected. Repeating this process many times with parameter sets randomly drawn from the prior distributions builds up a set of accepted parameters, which approximates the posterior parameter distribution ^10^.

We used a version of ABC based on sequential Monte Carlo sampling (ABC-SMC) ^12^. This reduces the required number of simulations by running multiple generations of simulations, with new generations of parameters based on the successful parameters from the previous generation. The threshold distance for parameter acceptance is reduced with each generation so that the distribution of accepted parameter sets increasingly approaches the target posterior distribution ^12^. We used the ABC-SMC implementation from the python package PyABC ^13^, and ran 15 generations using a population of 1000 particles (the required number of accepted parameter sets per generation).

#### 1.4. Parameter fitting results

Firstly, we see that the YFP clones in the *Notch1^+/+^* mice and the wild type intensity clones in the clonally induced *Notch1^+/-^* and *Notch1^-/-^* mice have inferred fitness values closely centered on neutral (**Extended data Fig. 4b, c, Table 1**). Both *Notch1^+/-^* and *Notch1^-/-^* clones are fitter than wild type clones, with *Notch1^-/-^* clones substantially fitter than *Notch1^+/-^* clones. Should this be paralleled in humans, this large increase in fitness conveyed by the second *Notch1* mutation may explain the strong selection and high prevalence of “double hit” *NOTCH1* mutant clones in aged human esophagus ^14^ (**Fig. 1c**).

**Table 1.** Inferred fitness parameter values from ABC fitting to lineage tracing data.

|  | Median inferred fitness | 95% credible interval |  |
| --- | --- | --- | --- |
|  |  | Lower | Upper |
| +/+ | 0.99 | 0.96 | 1.03 |
| +/+ in +/flox | 0.96 | 0.93 | 0.99 |
| +/+ in flox/flox | 1.03 | 0.96 | 1.12 |
| +/- | 2.3 | 2.0 | 2.6 |
| -/- | 7.0 | 6.2 | 8.6 |

In the cases of neutral clones (*Notch1^+/+^* and wild type intensity clones from induced mutant mice) the induction proportion was not constrained by the fitting (**Extended data Fig. 4b, Table 2**), because these clones have the same fitness as the surrounding wild type cells and therefore do not impact the growth of any other clones. The inferred induction proportion for the *Notch1^+/-^* clones was larger than for the *Notch1^-/-^* clones (**Extended data Fig. 4b, Table 2**). In simulations of the best fitting *Notch1^+/-^* parameters, the mutant clones were colliding at the later time points, reducing the clone sizes (**Video 1**). This high density of clones and extensive clonal collision was not occurring to such an extent in the experimental data (**Extended data Fig. 3g**), suggesting that the simple model is not fully capturing all details of the *Notch1^+/-^* clonal dynamics and may be overestimating the fitness of large *Notch1^+/-^* clones. However, the conclusions of the inferred fitness comparison are still valid: that the *Notch1^+/-^* clones have a clear growth advantage over wild type cells, and that the *Notch1^-/-^* clones have much larger growth advantage than the *Notch1^+/-^* clones.

**Table 2.** Inferred induction proportion parameter values from ABC fitting to lineage tracing data.

|  | Median inferred induction | 95% credible interval |  |
| --- | --- | --- | --- |
|  |  | Lower | Upper |
| +/+ | 0.054 | 0.002 | 0.099 |
| +/+ in +/flox | 0.051 | 0.004 | 0.098 |
| +/+ in flox/flox | 0.052 | 0.003 | 0.097 |
| +/- | 0.023 | 0.017 | 0.029 |
| -/- | 0.0045 | 0.0001 | 0.0085 |

#### 1.5. Simulations of haploinsufficiency

It is clear from the lineage tracing experiments and the model fitting results that *Notch1* is haploinsufficient, i.e. the heterozygous mutation gives a growth advantage to the clone. This growth advantage of the heterozygous mutant increases the chance of a double Notch1 mutant occurring because a higher proportion of the tissue becomes primed with the first mutation.

To illustrate the large difference haploinsufficiency makes, we ran simulations of haplo*insufficient* and haplo*sufficient* versions of *Notch1.* In the haplosufficient simulations, the first *Notch1* mutation has the same fitness as wild type cells, and a second *Notch1* mutation in the same cell increases the fitness to the best fitting *Notch-^-/-^* fitness (see **section 1.4**). In the haploinsufficient simulations, the first *Notch1* mutation provides a fitness advantage equal to the best fitting *Notch1^+/-^* fitness, then a second *Notch1* mutation in the same cell increases the fitness to the best fitting *Notch1^-/-^* fitness

We ran simulations with a very low mutation rate to parallel the experiments in which we age wild type mice and observed the appearance of *Notch1* mutant clones (**Extended data Fig. 4e, f, Video 2**). Each cell has a 0.0005% chance of gaining a mutation in each generation, with the mutation rate picked so that the haplosufficient simulations approximate the experimental data (**Extended data Fig. 4e**). In the simulations, a mutation can either appear on allele 1 or allele 2 of Notch1 with 50% chance of each. A cell with a single mutant allele has the heterozygous fitness of that simulation type (see above), a cell with both alleles mutated has the homozygous fitness. A second mutation to the same allele does not change the cell fitness. We ran the simulations for 5000 days or until all cells in the tissue had both Notch1 alleles mutated.

The simulations showed that it would take substantially longer for the tissue to be swept by the homozygous mutant clones if *Notch1* was haplosufficient (**Extended data Fig. 4e, f, Video 2**). This shows how the haploinsufficiency of *Notch1* is a key property that allows the gene to dominate clonal competition in normal EE.

### 2. DNA sequencing and coverage metrics

#### Custom baitset design

For targeted sequencing experiments, we used three different Agilent SureSelect custom bait capture kits. Lists of the genes included in each custom bait capture kits are detailed in **Supplementary table 1.** For the Human dataset, we used a kit comprising 322 genes designed to include frequently mutated genes in cancer. For the targeted sequencing of mouse tissue, we used a kit comprising 73 cancer related and Notch pathway related genes. This kit (baitset_3181471_Covered.bed) also includes several regions of chromosome 2, which carries *Notch1* gene. These regions, identified in a preliminary assay, allowed detection of SNP signal along the chromosome resulting in robust CNLOH analysis.

#### Targeted sequencing

For targeted sequencing, gridded tissue and tumor samples (**Methods**) were submitted at standard input of 200 ng of DNA for library preparation. Human micro-dissected samples and mouse subclonal samples were submitted for DNA library preparation allowing a reduced DNA input from 100-1000 cells. Samples were multiplexed and sequenced on Illumina HiSeq 2000 sequencer using paired-end 75-base pair (bp) reads.

For gridded tissue and tumor mouse samples, the average coverage across all genes after removing off-target reads, PCR duplicates were: DEN-SOR *Notch1^+/+^* grids: 478x; DEN-SOR tumors: 768x.

For the samples submitted at low DNA input, the average coverage across all genes after removing off-target reads, PCR duplicates were: Human biopsies: 33x; *Notch1^+/-^* subclonal biopsies: 194x.

### 3. Alignment, variant calling and gene selection analysis

#### Alignment

Paired-end reads were aligned with BWA-MEM (v.0.7.17, https://github.com/lh3/bwa) ^15^ with optical and PCR duplicates marked using Biobambam2 (v.2.0.86, https://gitlab.com/german.tischler/biobambam2). Human samples were aligned to the GRCh37d5 reference genome whilst Mouse samples were mapped to the GRCm38 reference.

#### -Mutation calling in the datasets

Methods used for mutation calling were adapted to the dataset, depending on if the samples were clonal/near-clonal or if we aimed at identifying somatic mutations present in a small fraction of cells within the samples.

Human micro-dissected samples and mouse subclonal samples for clonal or nearly clonal (**Fig. 1, Extended data Fig. 1, Fig. 3, Extended data fig. 5c-j, Supplementary tables 2, 4)**. Substitution mutations were called using the CaVEMan (Cancer Variants through Expectation Maximization, version 1.13.14) variant caller (http://cancerit.github.io/CaVEMan) ^16^. Insertions and deletions were called using cgpPindel (http://cancerit.github.io/cgpPindel, version 3.3.0) ^17^. Mutations were annotated using VAGrENT (https://github.com/cancerit/VAGrENT, version 3.3.3) ^18^. Mutations occurring outside of the targeted genes were removed by retaining only mutations annotated with the effects ‘missense’, ‘nonsense’, ‘ess_splice’, ‘frameshift’, ‘inframe’ or ‘silent’. The ShearwaterML pipeline (details below) was also used to analyze the Human dataset in order to report additional calling to the calls identified with CaVEMan and cgpPindel and allow a more exhaustive analysis of *NOTCH1* mutations (**Supplementary table 2**).

Gridded normal tissue and tumors (**Fig. 6, Extended data Fig. 7a,b, Supplementary table 7**) were sampled to analyze gene selection, map somatic mutations clones spanning multiple samples or present in a small fraction of cells within the samples. To do so, we used ShearwaterML algorithm from the deepSNV package (v1.21.3, https://github.com/gerstung-lab/deepSNV) to call for mutation events on ultra-deep targeted data ^14,19,20^. Instead of using a single-matched normal sample, the ShearwaterML algorithm uses a collection of deeply-sequenced normal samples as a reference for variant calling that enables the identification of mutations at very low allele frequencies.

The total coverage provided by the references used for each bait kits were as following. For the specifically designed Notch bait kit **(Supplementary table 1**), a total of 34 germline samples from the ears of the analyzed and the ears of extra mice providing a total coverage of 22885x. For the kit used for the Human sequencing, we used 68 germline samples for a total coverage of 14230x.

#### Gene selection (dNdS)

We used the maximum-likelihood implementation of the dNdScv algorithm (v0.0.1.0, https://github.com/im3sanger/dndscv) to identify genes under positive selection ^21^. dNdScv estimates the ratio of non-synonymous to synonymous mutations across genes, controlling for the sequence composition of the gene and the mutational signatures, using trinucleotide context-dependent substitution matrices to avoid common mutation biases affecting dN/dS. Values of dN/dS significantly higher than 1 indicate an excess of non-synonymous mutations in that particular gene and therefore imply positive selection, whereas dN/dS values significantly lower than 1 suggest negative selection.

#### Mutant tissue coverage

The size of mutant clones within each sample can be calculated by taking into account the area of the biopsy and the fraction of cells carrying a mutation within the sample, as described previously ^14,20^. The lower (=VAF) and upper (=2xVAF) bound estimates of the percentage of epithelium covered by clones carrying non-synonymous mutations in a given gene was calculated for each biopsy. The fraction of epithelium covered by the mutant genes was then calculated from the mean of summed VAF (capped at 1.0) of all the biopsies in the same tissue.

#### Merging of clones

For analysis of the mutational landscape in the normal epithelium from highly mutagenized mice, we performed gridding of the tissue into 2 mm^2^ biopsies and kept record of the relative positions of the samples (**Fig. 6b,c, Extended data Fig.7a).** In order to avoid counting the same mutation multiple times and to obtain a more accurate estimate of clone sizes clonal mutations that spanned between two or more adjacent biopsies were merged as individual events ^14,20^. To do this we calculated the mean number of shared mutations between biopsies at increasing distances, since the immediately adjacent samples are predicted to have more shared mutations than distant samples. Mutations common between samples closer than 3 mm were merged.

### 4. Copy number variation and copy neutral loss of heterozygosity

The *NOTCH1* locus is frequently affected by copy neutral loss of heterozygosity (CNLOH) events ^14,20^. CNLOH consists of the presence of two copies of either the maternal or paternal chromosome copy and none of the other. In sequencing data, this manifests itself as a change in the allele frequency of germline heterozygous SNPs (typically referred to as b-allele frequency, BAF), whilst the total coverage at the CNLOH event locus remains normal. The detection of a significant change in BAF and the absence of a change in coverage therefore constitutes CNLOH. To detect these events, we applied two pipelines based on previously published methods: One to detect alterations in coverage data and one for germline heterozygous SNPs.

Whilst a change in total coverage can readily be detected in data from both human and mouse samples, typically, inbred mouse strains do not contain enough germline heterozygous SNPs for the purposes of detecting a BAF imbalance. This project, however, contains data sets from mice of mixed-strain background where we did detect the presence germline heterozygous SNPs. We reasoned that, because the mouse strains used in this project are common, any reliable SNP call from the Mouse Genome Project ^22^, which is an effort to catalogue common strains, could be a viable germline heterozygous SNP candidate. Such a SNP would result from the mixture of strains, or rarely due to a *de novo* germline mutation. We therefore only considered SNPs that were reported by the Mouse

Genome Project and subsequently applied stringent coverage criteria to identify heterozygous SNPs in the matched control samples during the analysis (which is detailed further below).

Since both coverage data and germline heterozygous SNPs are available for both species, we applied both pipelines to all human and mouse samples.

#### 4.1. Detection of changes in total copy number

To call changes in total copy number in both human and mouse data we adapted QDNAseq for our purposes ^23^. Briefly, QDNAseq typically collects read counts in bins across the genome, normalizes the counts to produce relative coverage (commonly referred to as logR; coverage log ratio), adjusts coverage for GC content correlated artifacts, segments the data in sections of constant signal and finally calls regions that are significantly different from normal as either a gain or a loss.

The standard QDNAseq pipeline was adapted in three ways. First, since the datasets described in this manuscript always include a matched control sample, we included an extra step in the pipeline to adjust for coverage variability observed in the control sample to further reduce noise, as described in^24^.

Second, the calling of significantly different regions was adapted with a more conservative test, with the aim to produce robust calls. This step takes as input the segmented genome and each segment is tested by performing two one-sided t-tests: One to test for a gain and one to test for a loss. The test is performed between two distributions, one with the mean and standard deviation calculated from the logR of the segment (H1) and one with a mean of 0 (normal, unaltered logR) and the standard deviation of the segment (H0). This test may detect very small deviations for large segments, in-line with our aim for robust calls, we calculate the mean and standard deviation based on 100 sampled data bins for segments where more than 100 bins are available. The resulting p-values are adjusted for multiple testing (Bonferroni) and 0.05 is used as significance cutoff.

Finally, we included a pre-processing step to allow QDNAseq to run on targeted sequencing datasets. Typically, targeted sequencing data includes read-pairs that fall outside of the targeted genomic areas (referred to as off-target reads) and these off-target reads effectively form a shallow whole genome sequencing sample that can be used to call alterations ^25^. To obtain this shallow genome we first count reads in 1kb bins and subsequently remove any bin that overlaps with the targeted genomic coordinates. The remaining 1kb bins are then mapped to 1mb-sized bins, after which the regular pipeline is applied.

#### 4.2. Detection of allelic imbalance

The pipeline to detect allelic imbalance in both human and mouse data through germline heterozygous SNPs builds on previously published methods ^14,24^. This pipeline consists of 4 steps: (1) count reads for a set of reference SNPs (human: 1000 Genomes Project ^26^ mouse: Mouse Genomes Project) and determine which SNPs are heterozygous based on the matched control sample. (2) Reconstruct haplotypes to improve the accuracy of the b-allele frequencies as is used in the Battenberg algorithm ^24^, this step is skipped for mouse samples due to the requirement of reference haplotype data. (3) Segmentation of the data in regions of constant signal. (4) Calling of segments with significantly different signal and multiple-testing correction.

Allele counts are obtained for each potential SNP. For a read to be included in the count both the mapping quality and the base quality at the position within the read have to meet the below thresholds. A single read is counted for fragments where read pairs overlap, selecting the highest quality read to be included. These thresholds are applied to all counts in both the samples of interest and the matched controls.

| <u>Description</u> | <u>Species</u> | <u>Value</u> |
| --- | --- | --- |
| Minimum base quality | Both | 20 |
| Minimum mapping quality | Both | 35 |

A SNP was considered to be heterozygous when it met the criteria listed below in the matched germline sample. These thresholds aim to select bona fide heterozygous SNPs. Lower coverage thresholds were applied to the human samples due to their lower effective sequencing coverage. To compensate for the lower certainty in these samples, we subsequently increased the penalty of starting a new segment (segmentation Gamma).

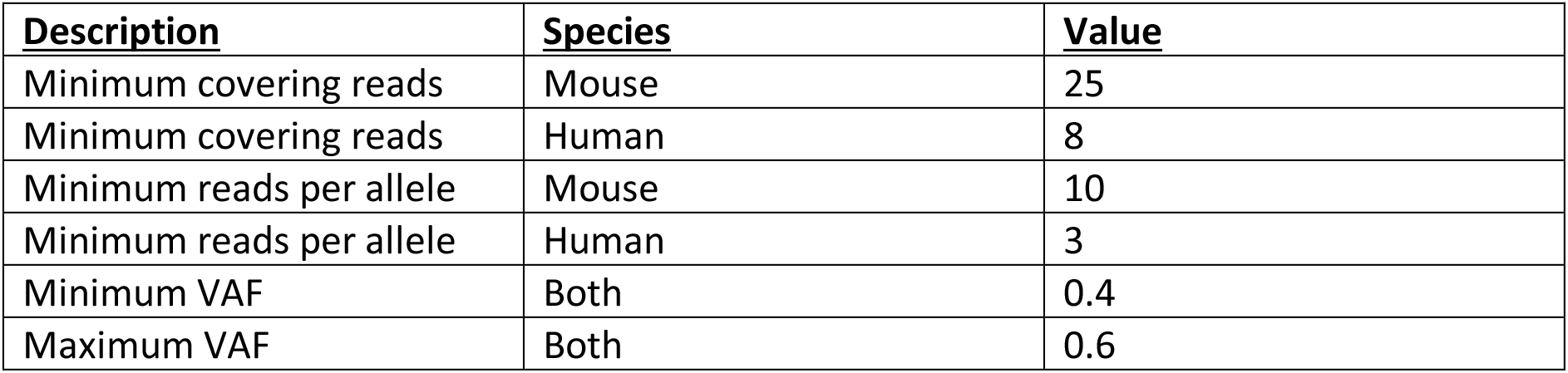

| <u>Description</u> | <u>Species</u> | <u>Value</u> |
| --- | --- | --- |
| Minimum covering reads | Mouse | 25 |
| Minimum covering reads | Human | 8 |
| Minimum reads per allele | Mouse | 10 |
| Minimum reads per allele | Human | 3 |
| Minimum VAF | Both | 0.4 |
| Maximum VAF | Both | 0.6 |

For human samples we next perform haplotype reconstruction to refine the BAF values, as is implemented in the Battenberg algorithm. Segmentation is subsequently performed via piecewise constant fitting, as implemented in the ASCAT software package ^24^, using the parameters listed below. This step results in segments that each have the following values: genomic start and end position, mean BAF in the sample of interest and mean BAF in the matched control.

| <u>Description</u> | <u>Species</u> | <u>Value</u> |
| --- | --- | --- |
| Gamma (reluctance to start a new segment) | Mouse | 6 |
| Gamma | Human | 10 |
| Kmin (minimum SNPs per segment) | Both | 3 |

Finally, for each segment the BAF of the sample of interested is tested against the BAF of the matched control sample. The BAF is required to deviate 0.1 from 0.5 (i.e. BAF < 0.4 or BAF > 0.6) before a segment is tested, with the aim to prohibit noise from generating low quality calls. Testing is subsequently performed via a two-tailed t-test where the BAF of all heterozygous SNPs in the sample of interest are tested against the BAF of the same SNPs in the matched control. The resulting p-values are adjusted for multiple-testing (Bonferroni). A segment is considered having a BAF imbalance when the adjusted p-value from this test is below 0.05.

| <u>Description</u> | <u>Species</u> | <u>Value</u> |
| --- | --- | --- |
| Minimum BAF deviation | Both | 0.1 |
| T-test significance threshold | Both | 0.05 |

### 5. Single-cell RNA sequencing analysis

#### 5.1. Alignment and transcript quantification

Alignment of the sequencing reads and expression quantification was performed for each library individually using the CellRanger pipeline version 3.0.2 (10xGenomics). We subsequently used EmptyDrops version 1.2.2 ^27^ to detect empty droplets in the raw feature count matrix output from CellRanger and discarded any barcode identified as an empty droplet. All the subsequent analysis described below was performed in R version 4.0.3 (https://www.R-project.org/) using the Seurat software package version 3.2.0 ^28^.

#### 5.2. Cell type detection

The pipeline to assign cell types first applies a number of filters per library to select healthy cells and remove any potential doublets. We filter on mitochondrial (MT) and overall expression levels: Cells are kept if their proportion of expression from MT genes is between 0.03 and 0.10, total expressed genes is between 2,500 and 6,500 and total expression is below 55,000 total unique molecule identifiers (UMI). Genes are kept if they are expressed in 30 cells or more per library.

After filtering, expression is normalized and adjusted for proportion of MT expression, total expression, total number of expressed genes and predicted cell cycle state using the SCTransform function from Seurat. So far, all steps have been applied to the four libraries individually. The libraries are next combined using Seurat’s SCT integration methodology that aims to incorporate libraries, taking into account possible batch effects. To capture the majority of the variation we required 3,000 genes to be used for the integration. Next, PCA was performed and we selected the top 30 principal components to continue into UMAP dimensionality reduction, clustering of cells and detection of variable genes between clusters. A plot of the UMAP space revealed good mixing of the four libraries throughout (**Fig. 4g**).

Cell types were identified through a series of marker genes. For keratinocytes we used *Krt14*, *Tgm3*, and *Lor* and required a per cell cluster average of at least 50, 5 and 10 transcripts per million (TPM) respectively (**Extended data Fig. 6e**). For fibroblasts we used *Col1a1* with an average cluster TPM cut- off of greater than 20, for endothelial cells we used *Pecam1* requiring an average cluster TPM over 2 and, finally, for immune cells we used *Cd83*, *Cd84*, *Cd86*, *Cd52* and *Ptprc* with average cluster TPM values of 1, 1, 0.4, 50 and 0.25 respectively. **Extended data Fig. 6c** shows the expression of a number of selected markers overlaied on the UMAP space. **Extended data Fig. 6e** shows expression of a number of keratinocyte marker genes.

Visual inspection of the assignment through a UMAP plot revealed that, even though the cell types were well separated (**Fig. 4h, Supplementary table 6**), 35 cells had been marked as keratinocytes, but visually clustered with the other cell types. These cells were flagged and their cell type label was set to NA to reflect their ambiguous status. These cells were not used for further analysis.

The number of cells per cell type, per library were counted and divided by the total number of cells per library to produce the data shown in **Fig. 4h**.

#### 5.3. Analysis of keratinocytes

Having identified which cells are keratinocytes, we restarted the analysis from the raw UMI counts, but now only including those cells that were previously marked as keratinocytes. The analysis differed from that above in two critical ways: 1. No cell cycle adjustment was performed as the cell cycle state and keratinocyte differentiation are deeply linked, with basal cells cycling and differentiating cells having exited the cell cycle, and 2. as now only keratinocytes were included, we increased the minimum number of genes expressed per cell to 3,000 to strictly select for highly viable cells.

After integration of the four libraries, we performed a PCA and selected the top 10 principal components for further analysis. Clustering was performed and significant differentially expressed genes were identified through standard Seurat functions. Cells were assigned to a cell cycle phase using the Seurat function CellCycleScoring (**Fig. 4j, Supplementary table 6**) and we subsequently counted the cells per phase, per library and divided these counts by the total number of keratinocytes in each library to produce the data shown in **Fig. 4j.**

To determine the number of basal cells per library we made use of the visual dividing line in the UMAP space marked by a clear drop in expression of *Krt14* (basal cell marker gene) and the emergence of that of differentiation markers *Krt4* and *Tgm3* (**Extended data Fig. 6e**). The line separating the ‘circl’ shape from the ‘tail’ shape was defined to roughly capture the area where this expression divide manifests itself. We subsequently counted the cells from each library on both sides of the line to obtain the data shown in **Fig. 4i (Supplementary table 6)**. We note that the exact placement of the dividing line matters little for the differences in proportions between libraries, because of the good mixing of cells from both wild-type and knock out libraries across the UMAP space (**Fig. 4g**).

---

**Extended data Figure 1:**
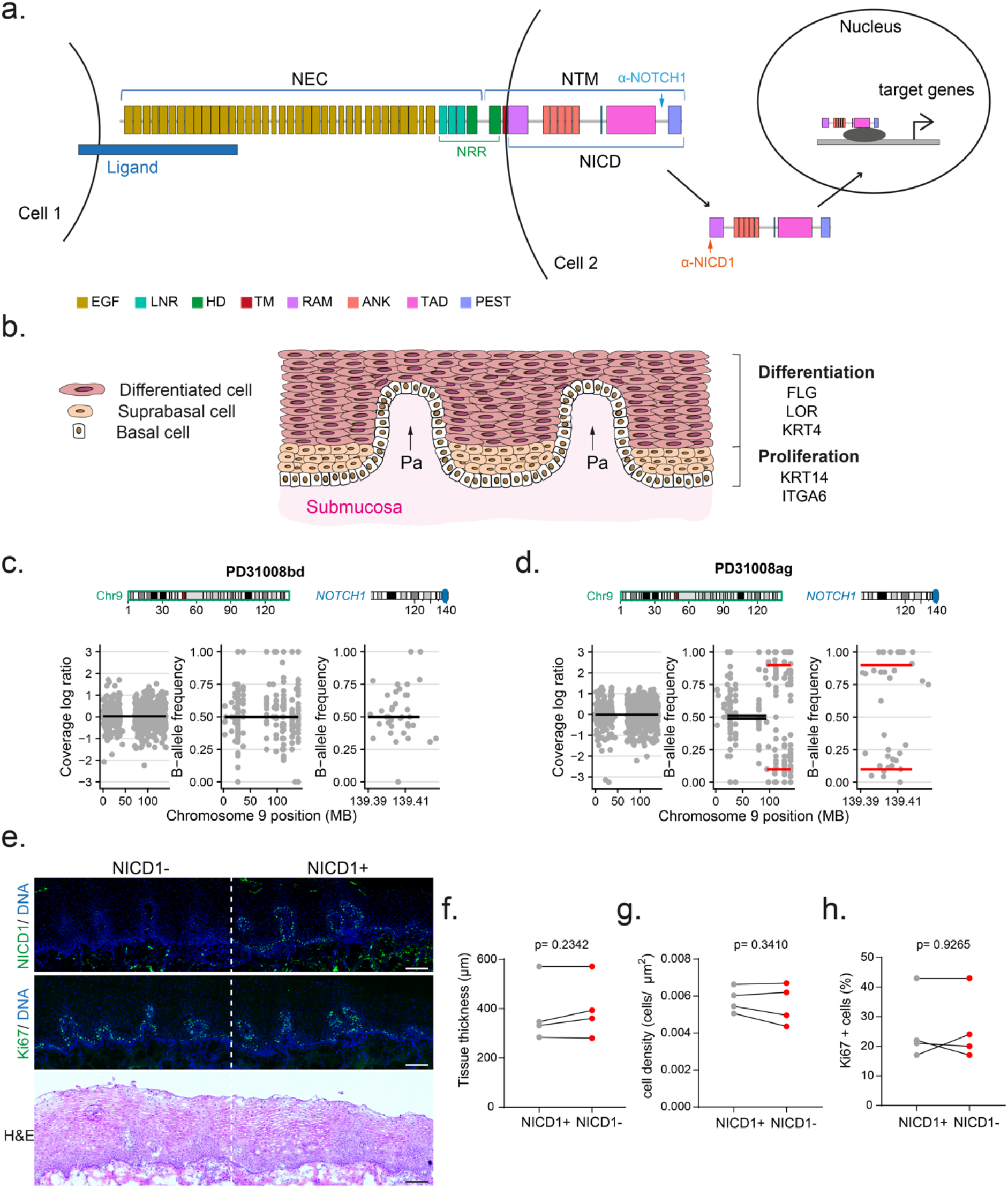
Aging human esophageal epithelium is colonized by *NOTCH1* mutant clones. **a.** Schematic showing Notch signaling and NOTCH1 subunits. NOTCH1 is a membrane receptor composed of an extra-cellular domain (NEC) and a transmembrane and cytoplasmic unit (NTM). Activation by ligand expressed on a neighboring cell leads to two successive cleavages releasing the intracellular domain (NICD) which translocates into the nucleus and interacts with partners to activate target genes. Functional domains of NOTCH1 are indicated, arrows show epitopes recognized by the antibodies used for anti-NOTCH1 (blue) and anti-NICD1 (orange) immunodetection **b.** Human esophageal epithelium. The basal layer expresses *ITGA6* and *KRT14*. Proliferating cells are found in the basal and lower suprabasal cell layers which express *KRT14*. Differentiating cells leave the proliferative compartment and migrate to the tissue surface, expressing the early and late differentiation markers *KRT4* and *LOR*, *FLG* respectively. Cells are continually shed from the epithelial surface and replaced by proliferation in the lower cell layers. Pa, papillae. **c, d**. Figures of copy number calls for the Human microbiopsies shown in **Fig.1b, c** and **Supplementary table 2c**. On the left plot, analysis of off-target reads show no difference in total copy number along chromosome 9 for both c and d. Middle and right hand plots show the B allele frequency (BAF) of germline heterozygous SNPs along chromosome 9 and the *NOTCH1* locus, respectively. Red lines denote a significant difference compared to a matched control. The figures in **d** show a high BAF for sample PD31008ag without a change in total copy number reflecting copy neutral loss of heterozygosity. **e.** Successive sections of esophagus from middle aged and elderly donors were stained for active NOTCH1 (NICD1), proliferation marker Ki67 and with Hematoxylin and eosin (H&E). Images are representative from transversal sections analyzed in 4 middle-age and elderly donors. Scale bars, 100µm. **f, g, h.** Tissue thickness (**f**), cell density (**g**) and proportion of proliferative cells (**h**) were measured in NICD1 positive and negative areas, on sections stained for NICD1, Ki67 and DAPI. Each dot represents a donor. For **f**, NICD1+, n=14 areas from 4 donors, NICD1-, n=15 areas from 4 donors. For **g**, NICD1+: 10795 cells from 4 donors, NICD1-: 11593 cells form 4 donors. For **h**, NICD1+: 5402 cells from 4 donors, NICD1-: 6204 cells form 4 donors. Two-tailed paired Student’s t-test. See **Supplementary tables 1-3.**

**Extended data figure 2:**
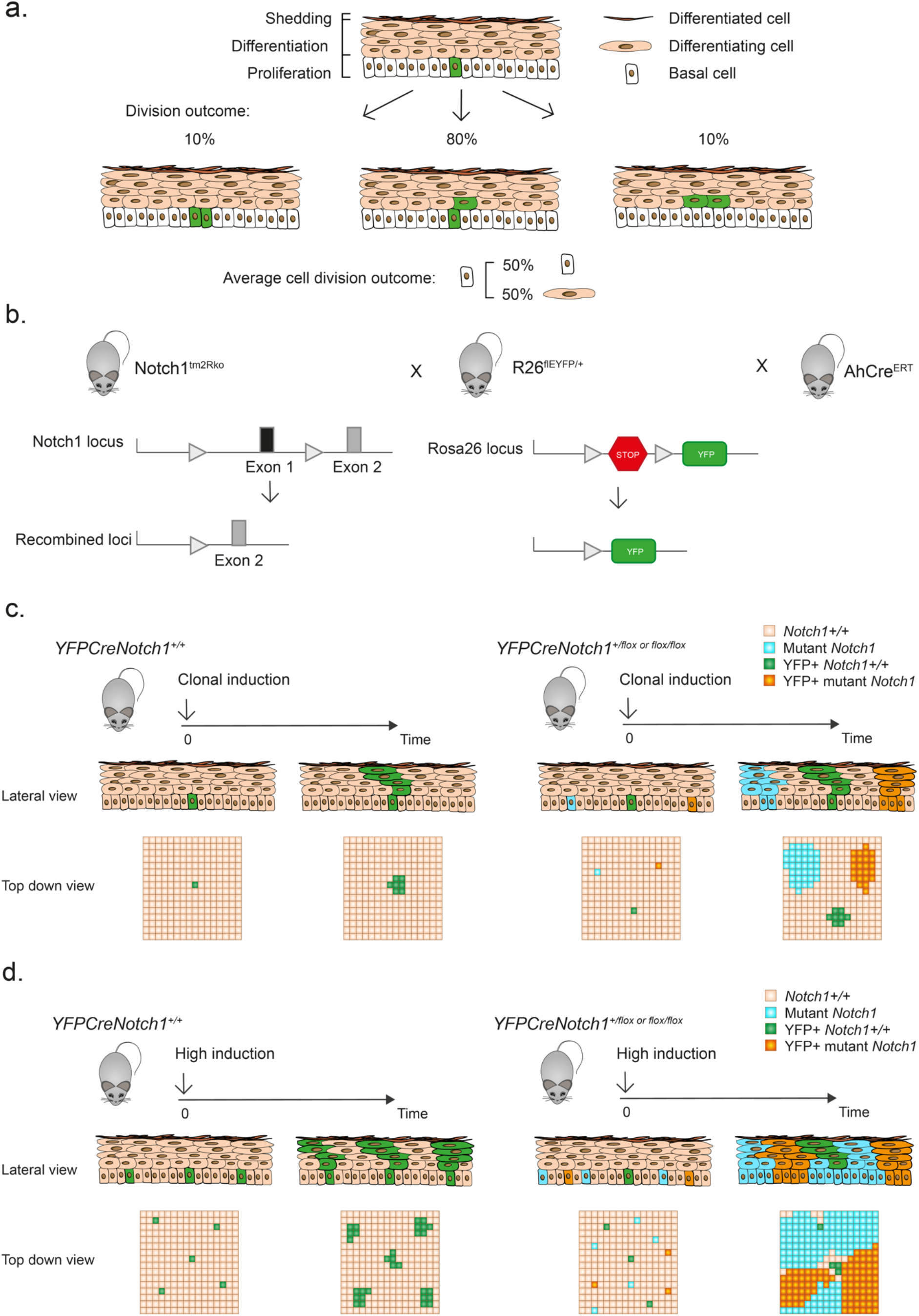
Lineage tra1cing of *Notch1* mutant cells in mouse oesophageal epithelium. **a.** Structure and cellular homeostasis in mouse esophageal epithelium. The basal layer contains progenitor cells that divide to generate progenitor and differentiating daughter cells. Differentiating basal layer cells exit the cell cycle and migrate into the suprabasal layers, moving towards the surface of the epithelium from which they are shed. The division of a progenitor cell (green) produces two progenitors, two differentiating cells or one cell of each type. In homeostatic tissue, the likelihood of each division outcome is balanced and gives on average 50% of progenitors and 50% of differentiating cells across the progenitor population. **b.** *YFPCreNotch1* conditional knock-out mouse strain. *LoxP* sites (grey arrows) flank exon1 of the *Notch1* gene. *Notch1* floxed animals were crossed with *Rosa26^floxedYFP^* mice carrying a conditional Yellow fluorescent protein (YFP) reporter targeted to the *Rosa26* locus and with *AhCre^ERT^* mice carrying an inducible *Cre* recombinase. **c.** For lineage tracing, triple mutant mice were treated with inducing drugs at a dose that resulted in recombination of *Notch1 (blue)*, expression of YFP (green) or both (orange) in scattered individual esophageal basal cells (clonal induction). The recombined cells may expand into clones detected by the reduced intensity (+/-) or absence of NOTCH1 (-/-) and expression of YFP detected by immunostaining. Samples were collected at different time points after induction and the number and location of cells in each clone determined by 3D confocal imaging of sheets of epithelium. **d**. Triple mutant mice were induced with a high dose of drugs, allowing recombination of cells at high density in the tissue. In the case of mutant clones with a competitive advantage over wild type cells, this protocol allowed the coverage of the tissue by mutant clones relatively shortly after induction.

**Extended data Figure 3:**
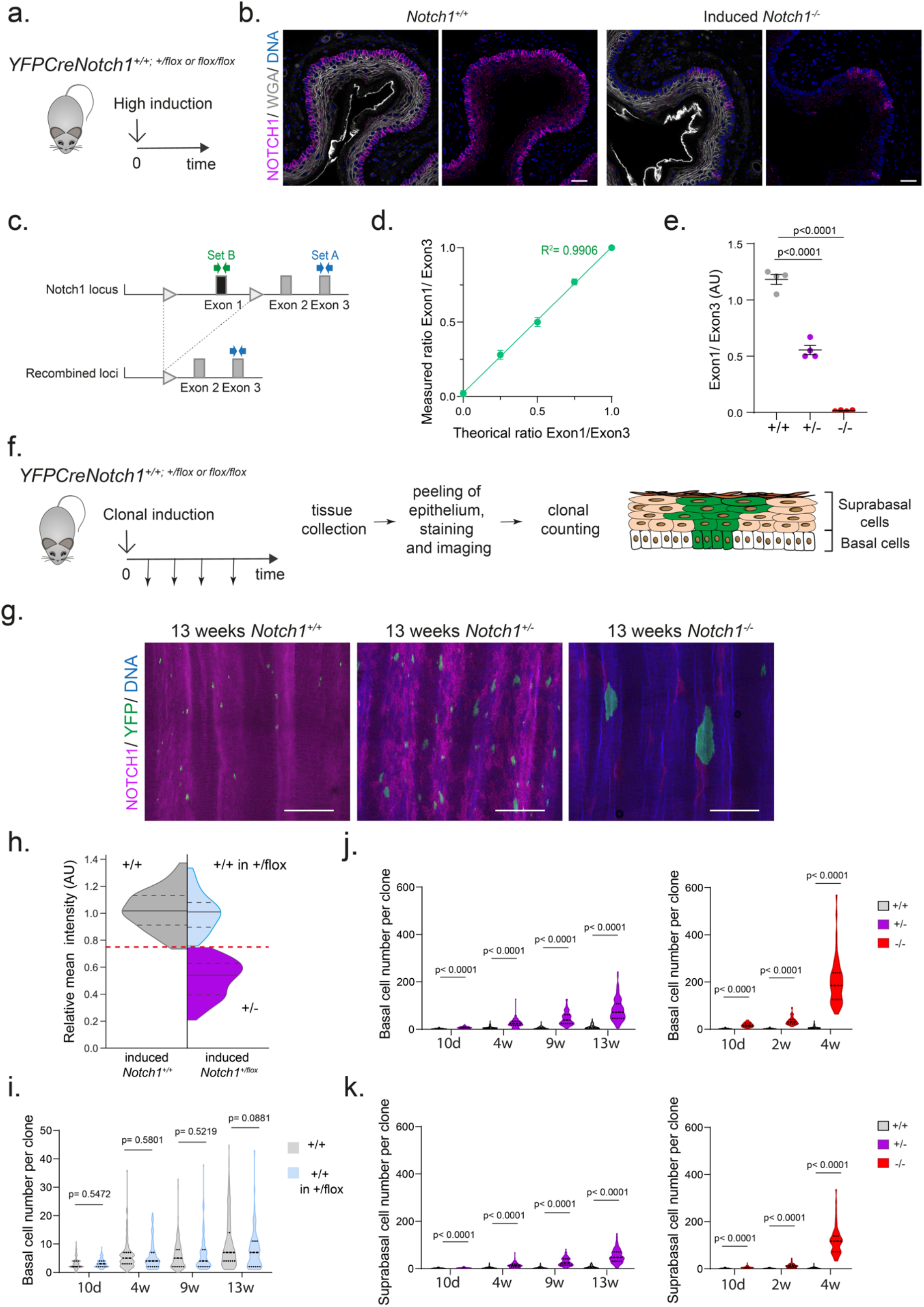
Dynamics of *Notch1*^+/-^ and *Notch1^-/-^* clones. **a.** Protocol for **b-e**. *YFPCreNotch1^flox/flox^* or *^+/flox^* mice were induced to give a high level of recombination and aged to allow Notch1 mutant cells to colonize the epithelium. Non-induced *YFPCreNotch1^flox/flox^* mice were used as wild type controls (+/+). Esophageal epithelium was then collected. **b.** Immunostaining of transversal section of esophageal tissue 10 days after high induction. NOTCH1 (magenta) expression is lost in most basal cells in induced *Notch1^flox/flox^* mice. Wheat germ agglutinin (WGA) marks the cell membrane (grey). Scale bars, 30µm. **c.** Quantitative PCR assay for *Notch1* recombination. Primer set B measures the loss of the floxed exon1 of Notch1 by Cre excision after induction protocol. Primer set A allows intragenic normalization within *Notch1* exon 3. **d.** Validation of the linearity of the recombination assay against a standard curve reproducing different recombination rates with Exon1/Exon3 ratio of 1, 0.75, 0.5, 0.25 and 0. AU, arbitrary units. **e.** Recombination assay was performed on the esophageal epithelia shown in **Fig.2a-c.** Exon1/Exon3 ratio is halved in induced *Notch1^+/-^* mice compared to wild type mice and null in induced *Notch1^-/-^* mice suggesting full recombination of the esophageal epithelium. Mean± SEM, each dot represents a mouse, n=4. One-way ANOVA; adjusted, p values from Tukey’s multiple comparisons test against wild type. **f**. Protocol for genetic lineage tracing. *YFPCreNotch1^flox/flox^* or *^+/flox^* or *^+/+^* mice were induced at clonal level, tissue was collected, tissue was peeled and stained by whole mount for NOTCH1, YFP and DAPI and cohorts of YFP+ clones imaged at different time points. Basal and suprabasal cells were counted in mutant or control clones categorized on the basis of NOTCH1 staining (**Extended data Fig.2a**). **g.** *Notch1^+/+^, Notch1^+/-^* and *Notch1^-/-^* clones 13 weeks post clonal level induction in esophageal epithelial wholemounts stained for NOTCH1 (magenta), YFP (green) and DNA (blue). Scale bars, 500µm. **h. d**. *YFPCreNotch1^+/+^* and *YFPCreNotch1^+/flox^* mice were induced at clonal level and intensity of NOTCH1 staining was measured in the basal layer of YFP+ clones. Recombination of YFP is independent of the *Notch1* locus so YFP clones may be *Notch1^+/+^* or *Notch1^+/-^* in *YFPCreNotch1^+/flox^* animals depending on whether *Notch1* was recombined. Intensity of clones in *YFPCreNotch1^+/+^* mice served as reference for discriminating *Notch1* wild type from *Notch1^+/-^* clones in the induced *Notch1^+/flox^* mice. Violin plot shows the distributions of relative NOTCH1 staining intensity of YFP+ clones in *Notch1^+/+^* mice (left, grey, n=3 mice) and *Notch1^+/-^* mice (right, n=7 mice). Black lines show median and quartiles. Red dashed line indicates the threshold relative intensity (75) determined based on the normal approximation of the distribution in wild type mice and the two-component Gaussian mixture model approximation of the distribution in induced *Notch1^+/flox^* mice. Clones with intensity >75 in induced *Notch1^+/flox^* mice are categorized as wild type (+/+ in +/flox, blue), the other clones are categorized as *Notch1^+/-^* (+/-, purple). **i.** Validation of clonal genotype from NOTCH1 immunostaining. Violin plots show the number of basal cells/clone in wild type (+/+) and wild-type intensity populations from induced *Notch1^+/-^* tissue (+/+ in +/flox). Lines show median and quartiles. Groups do not show statistical difference in clonal size at 10 days, 4, 9 and 13 weeks. Number of mice (clones) for +/+ and +/+ in +/flox respectively at 10 days: 3 (206)/ 5 (94); 4 weeks: 3 (143)/ 4 (78); 9 weeks: 3 (132)/ 4 (72); 13 weeks: 3 (126)/ 7 (65). Two tailed unpaired Student’s t-test against +/+ at each time point. **j, k.** Basal cell number (**j)** and suprabasal cell number (**k**) per clone is shown with violin plots at different times after induction. YFP positive *Notch1^+/-^* (left panel) or *Notch1^-/-^* (right panel) clonal cell counts are shown in comparison to *Notch1^+/+^* clones. Lines show median and quartiles. Fusion of expanding clones prevented quantification of clone size beyond 4 weeks for *Notch1^flox/flox^* tissue and beyond 13 weeks for *Notch1^+/flox^* tissue. Number of mice (clones) for +/+ at 10 days, 2 weeks, 4 weeks, 9 weeks, 13 weeks respectively: 3 (206)/ 3 (155)/ 3 (143)/ 3 (132)/ 3 (126). Number of mice (clones) for +/- at 10 days, 4 weeks, 9 weeks, 13 weeks respectively: 5 (84)/ 4 (97)/ 4 (68)/ 7 (107). Number of mice (clones) for -/- at 10 days, 2 weeks, 4 weeks respectively: 6 (68)/ 3 (69)/ 9 (63). P values from two-tailed unpaired Student’s t-tests against +/+ at each time point.

**Extended Data Figure 4:**
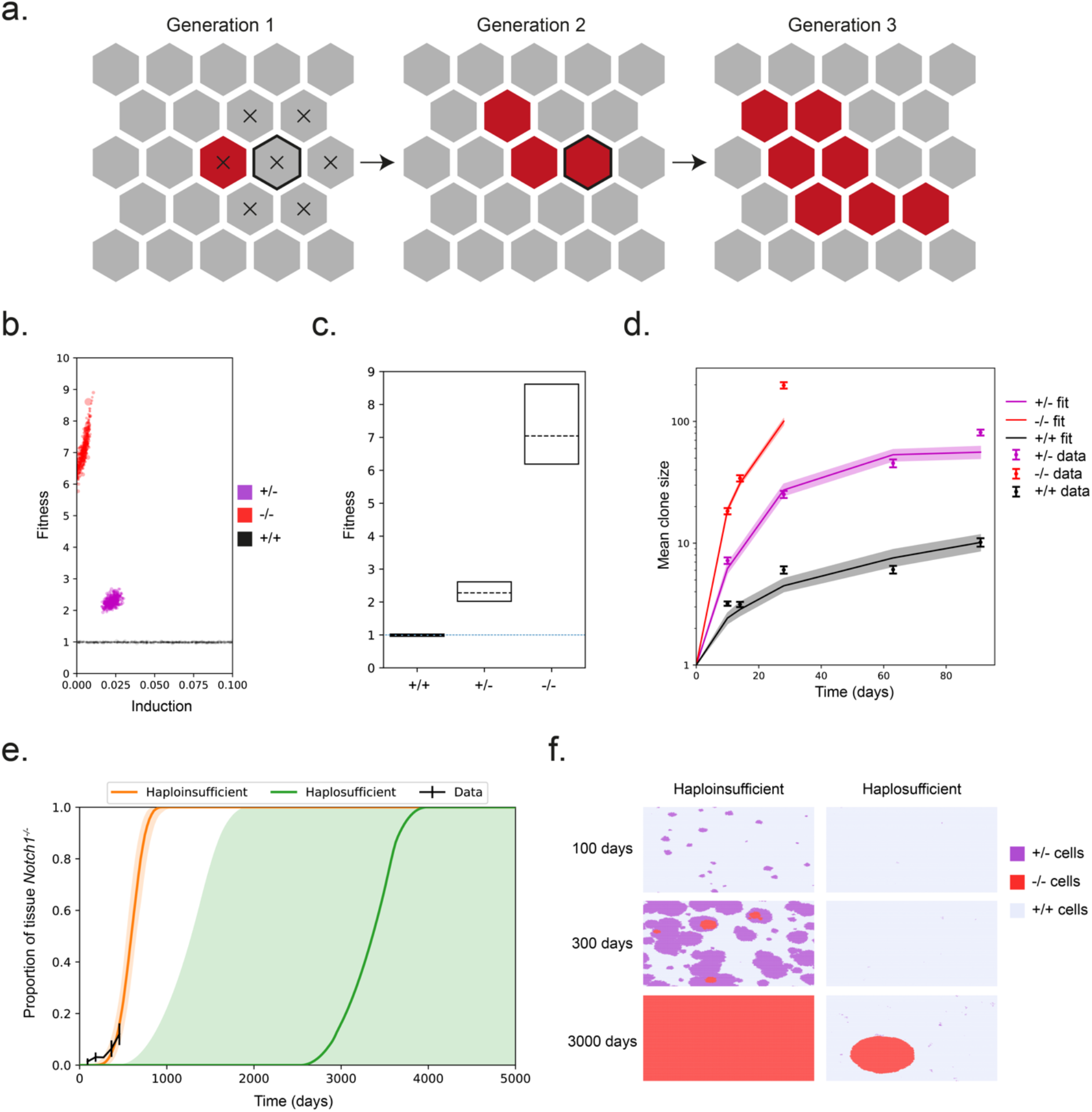
Modelling *Notch1* mutant clone expansion. **a.** 2-dimensional Wright-Fisher style model of clone dynamics in the esophagus. The basal layer is made up of a hexagonal grid of cells. At time zero, a small proportion of cells are assigned to be mutant (red) and the rest wild type (grey). Cells in the next generation are picked from neighboring cells – e.g. the cells which can be placed in the outlined position in generation 2 are those marked with an X in generation 1. Mutant cells with higher fitness have a higher probability of generating daughter cells in the next generation and hence can expand into large clones (**Supplementary Note**). **b.** Plot shows the inferred induction proportion and inferred fitness values from ABC fitting to the lineage tracing data (**Supplementary Note**) for *Notch1^-/-^* (red), *Notch1^+/-^* (purple) and *Notch1^+/+^* (black) animals. Each dot shows an “accepted” parameter set. **c.** Distributions of acceptable values of the fitness parameter. Dashed lines and box show median inferred fitness and 95% credible intervals. A fitness of 1 is neutral. **d.** Mean clone sizes from simulations of the parameters at the peak of the acceptable distributions (see **Supplementary note**). Median and 95% confidence intervals of 100 simulations shown for the simulation curves. Mean ± SEM are shown for the experimental data. **e.** Proportion of tissue covered by *Notch1^-/-^* clones over time in simulations using the best fit fitness parameter for *Notch1^-/-^* fitness. *Notch1^+/-^* is either assumed to be neutral (haplosufficient, green) or to have the best fitting fitness parameter to the experimental analysis of *Notch1^+/-^* clones (haploinsufficient, orange). Curves show median and shaded areas show 95% confidence intervals. **f.** Snapshot images at 100 days, 300 days and 3000 days of video modelling using the simulations shown in e. On the left, *Notch1^+/-^* cells are haploinsufficient (fitting to observed experimental data), on the right the *Notch1^+/-^* cells are assumed to be haplosufficient (neutral fitness). See **Supplementary Note**.

**Extended Data figure 5:**
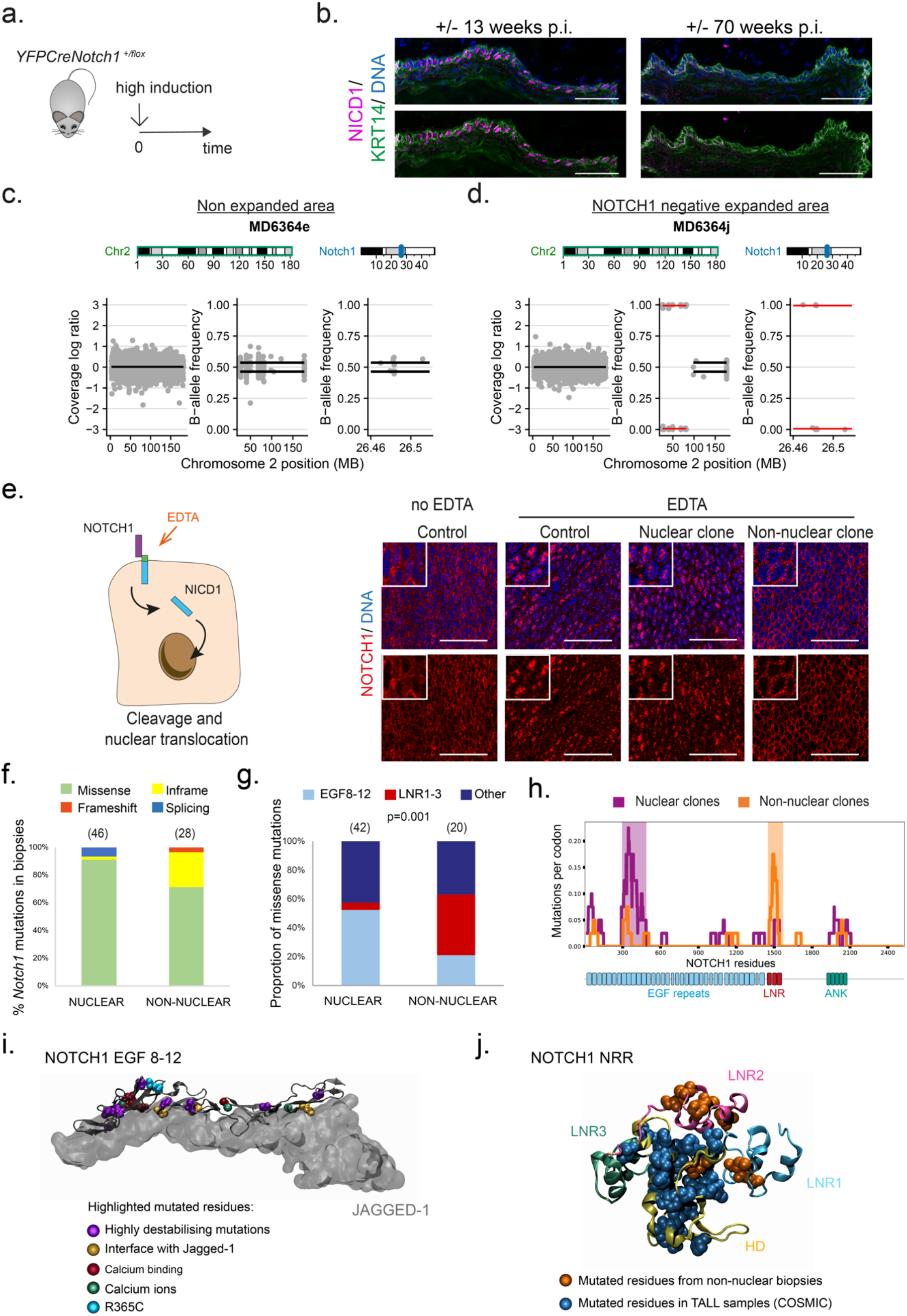
Analysis of spontaneous mutant clones in *Notch1^+/-^* aged esophageal epithelium. **a.** Protocol. *YFPCreNotch1 ^+/flox^* mice were induced at high density and aged up to 78 week-old. **b**. Representative NICD1 (magenta) and KRT14 (green) staining of sections of esophagi from *Notch1^+/-^* mice, 13 weeks and 70 weeks after induction. Images are representative examples from 3 and 5 mice respectively. Scale bars, 50µm. **c, d.** As in Fig. 3e, expanded and non-expanded areas were dissected and targeted sequencing performed. Plots show representative examples of copy neutral LOH analysis from non-expanded normal area (**c**) and NOTCH1 negative expanded area carrying CNLOH (**d**) Total copy number and loss of heterozygosity analysis from targeted sequencing. The left panel contains the read coverage of off-target reads across chromosome 2 and shows no changes in total copy number were detected. The middle and right panels show the B allele fraction (BAF) of germline heterozygous SNPs along chromosome 2 (middle) and at the *Notch1* locus (right). Significantly different values for total copy number or BAF imbalance are shown with red lines. The NOTCH1 negative sample MD6364j shows CNLOH at clonal level. **e**. Left panel. Aged *YFPCreNotch1^+/-^* mice esophagi were incubated in 5mM EDTA solution, activating NOTCH1 cleavage in the absence of ligand, stained for NOTCH1 (red), YFP and DNA (blue) and control and clonal areas were sampled and identified as shown in Fig. 3e-g. Right panel shows representative pictures of NOTCH1 positive areas. Non expanded normal areas show nuclear NOTCH1 only following EDTA treatment as opposed to non- EDTA treated cells displaying both nuclear and non-nuclear staining. EDTA treated positive clones display either nuclear staining (nuclear clone) or extra nuclear NOTCH1 (non-nuclear clone). Scale bars, 50 µm **f.** Analysis of the type of *Notch1* mutations in mutant clones with NOTCH1 nuclear staining (nuclear) and without nuclear staining (non-nuclear) imaged as in **e**. Number of samples is indicated in brackets. Redundant samples defined as biopsies sharing the same mutation and spaced <1mm away in the tissue were counted once. **g.** Location of missense mutations in nuclear and non-nuclear samples in NOTCH1 protein. Number of samples is indicated in brackets. P= 0.001, Chi-square test.**h.** Distribution of NOTCH1 codon alterations in missense mutants in nuclear (purple, n=42 mutations) and non-nuclear (orange, n=20) clones. Purple shadow highlight the cluster of mutations in EGF repeat 8-12, Orange shadow highlight the cluster of mutations at the LNR repeats. **i.** Missense mutations in *NOTCH1* EGF8-12 in samples with nuclear staining. Residues containing missense mutations are highlighted on the structure of rat Notch1 EGF8-12 bound to Jagged-1 (PDB 5UK5). Mutated residues involved in calcium binding are shown in dark red, residues on the interface with NOTCH1 ligands are shown in yellow, and residues with highly destabilizing mutations (FoldX ΔΔG > 2kcal/mol) are shown in purple. R365C, which does not fit into any of the previous categories is shown in blue. Calcium ions are shown in green, n=21 missense mutations. **j.** Mutated residues highlighted on the 3D human NRR structure (PDB 3ETO). Missense mutations in NOTCH1 NRR in biopsies with non-nuclear staining are shown in orange (n=9 mutations). Activating missense mutations from human T-cell acute lymphoblastic leukemia (T-ALL) samples from COSMIC (https://cancer.sanger.ac.uk/cosmic ^25^) are shown in blue (n=153 mutations). The proportion of missense mutations between LNR1-2 and LNR3- HD in T-ALL and non-nuclear NOTCH1 stained esophagus is significantly different (Two-sided Fisher exact tests, p=1.48e^-10^). LNR1, blue; LNR2, pink; LNR3, green; HD domains, yellow. See **Supplementary table 4**.

**Extended data Figure 6:**
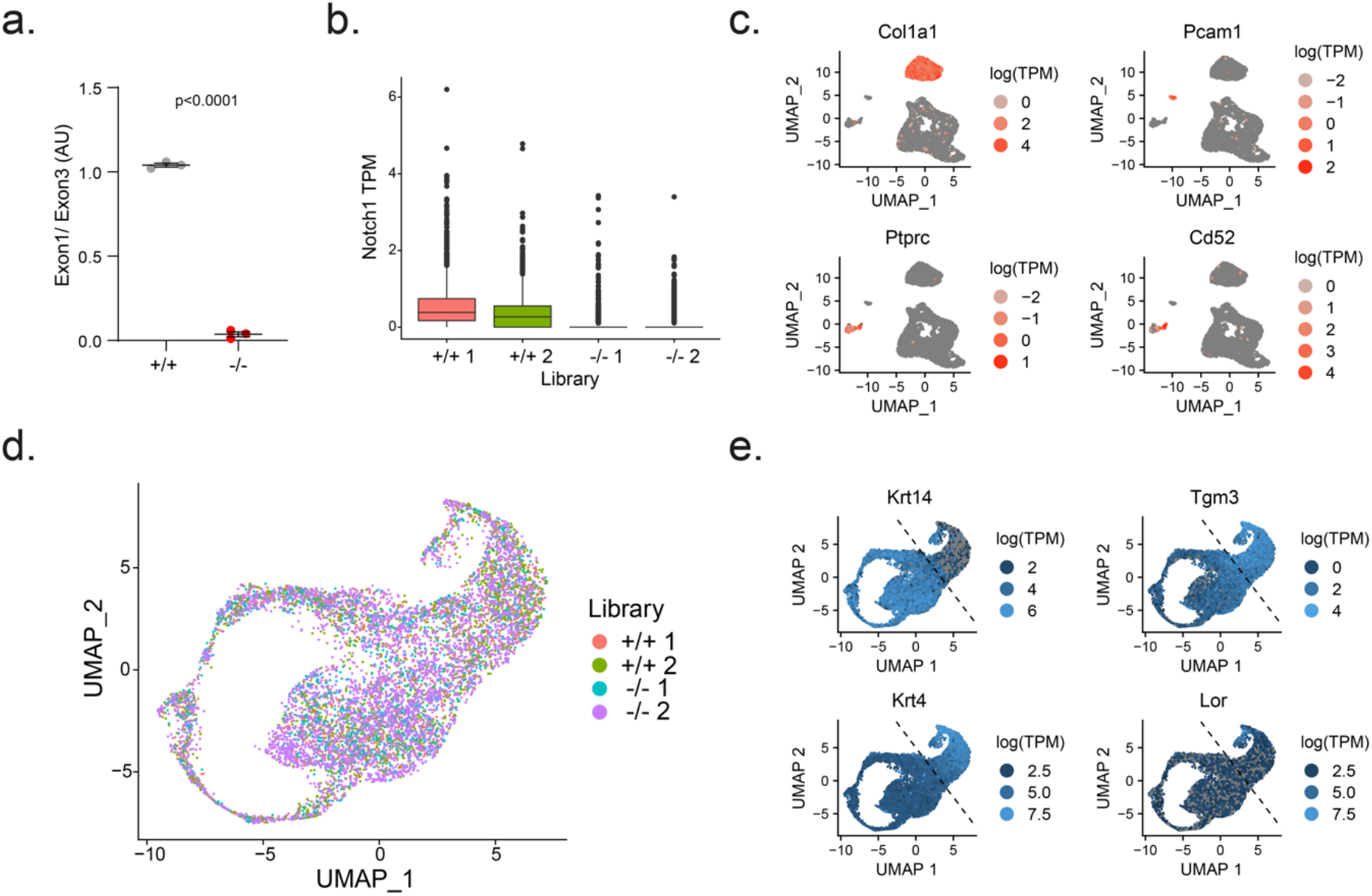
single cell RNAseq of *Notch1* mutant tissue. **a.** qPCR recombination assay of *Notch1* exon1 in peeled esophageal epithelium from *Notch1^flox/flox^* mice induced once at the same dose as the one used for the sc-RNAseq and collected 4 weeks after induction. Data is shown in comparison to wild type tissue from the same qPCR assay. Mean ± SEM, each dot represents a mouse, n=3 mice. Two-tailed unpaired Student’s t-test. **b**. Esophageal epithelium was processed for single cell RNAseq as shown in **Fig.4f**. Box plots show transcripts per million (TPM) of *Notch1* in the cells from each library (n=2 biological replicates per genotype). **c**. UMAP plots show some markers that were used for cell type annotations of fibroblasts (*Col1a1*, top left), endothelial cells (*Pcam1*, top right) and immune cells (*Ptprc*, bottom left; *Cd52*, bottom right). See **Supplementary Note**. **d.** UMAP plot shows an overlay of the cells annotated as keratinocytes from each library (+/+1, n=1555; +/+2, n=1931; -/-1, n=1403; -/-2, n=3919). **e.** UMAP plots show markers of keratinocyte differentiation highlighting basal cells (Krt14, top left), differentiating cells (*Tgm3*, top right, *Krt4*, bottom left) and cornified cells (*Lor*, bottom right). Dashed black line shows the threshold defining basal cells on the left and suprabasal cells on the right following *Krt14* and *Krt4* expression levels (**Supplementary Note**). AU, arbitrary unit. See **Supplementary table 6**.

**Extended Data Figure 7:**
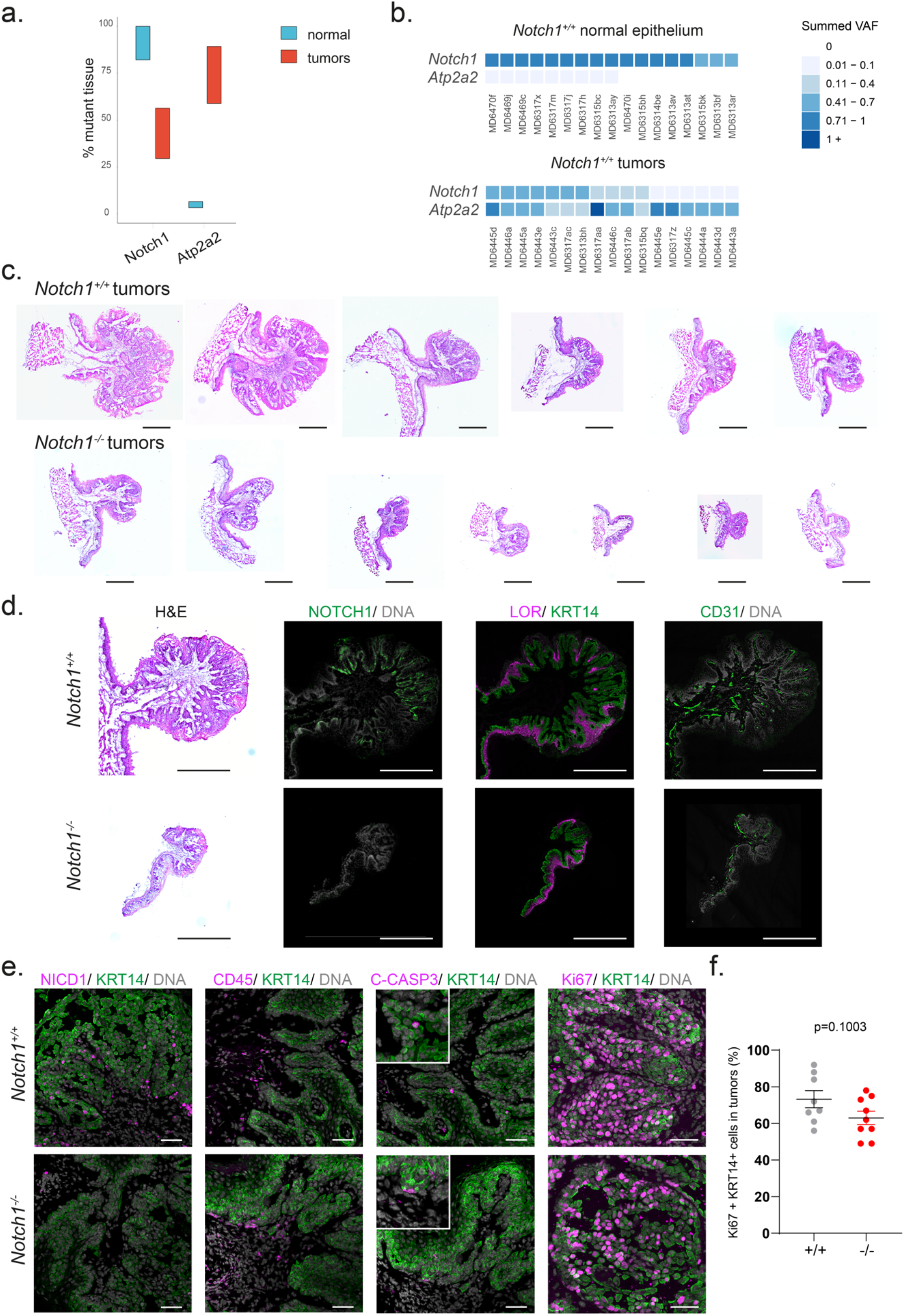
Genetic and histological characterization of mouse esophageal tumors. **a.** As in **Fig.6a**, mouse esophageal tissue and tumors from carcinogenesis assay were collected 28 weeks after treatment and processed for targeted sequencing. Proportion of tissue mutant for *Notch1* and *Atp2a2* in tumors and in normal epithelium. Estimates are from sum of VAFs of non-synonymous mutations for *Notch1* and *Atp2a2* (n= 115 normal epithelial biopsies from 6 mice, n=17 tumors from 7 mice; **Supplementary Note**). **b.** Summed VAF of protein altering mutations in the indicated genes is shown for randomly picked and representative normal gridded tissues (upper, n=17 out of 115 biopsies) and each 17 sequenced tumors (lower). **c.** Hematoxylin and eosin (H&E) staining of tumors from uninduced wild type and induced *Notch1^-/-^* mice treated with DEN and Sorafenib and aged for 28 weeks as shown in Fig. 6. Pictures are representative of n=19 wild type and n=11 *Notch1^-/-^* stained tumors. Scale bars, 500 μm. **d, e.** *Notch1^+/+^* and *Notch1^-/-^* tumors were sectioned and stained, from left to right, (**d**) for H&E, NOTCH1, differentiation markers (KRT14 and LOR), endothelial cell marker CD31, (**e**) active NOTCH1 (NICD1), immune cell marker CD45, for apoptosis marker cleaved Caspase 3 and proliferation marker Ki67. Images are representative of n=10 wild type tumors and n=11 *Notch1^-/-^* tumors for all immunostainings and cleaved Caspase 3 (n=4 tumors). Scale bars in **d**, 500 μm, in e, 50 μm. **f.** Percentage of Ki67 positive, KRT14+ keratinocytes within tumors from *Notch1^+/+^* and *Notch1^-/-^* esophagus. Mean ± SEM, each dot represents a tumor, n=8 tumors from 5 *Notch1^+/+^* mice and n= 9 tumors from 6 *Notch1^-/-^* mice. Two-tailed unpaired Student’s t-test.

**Extended data figure 8:**
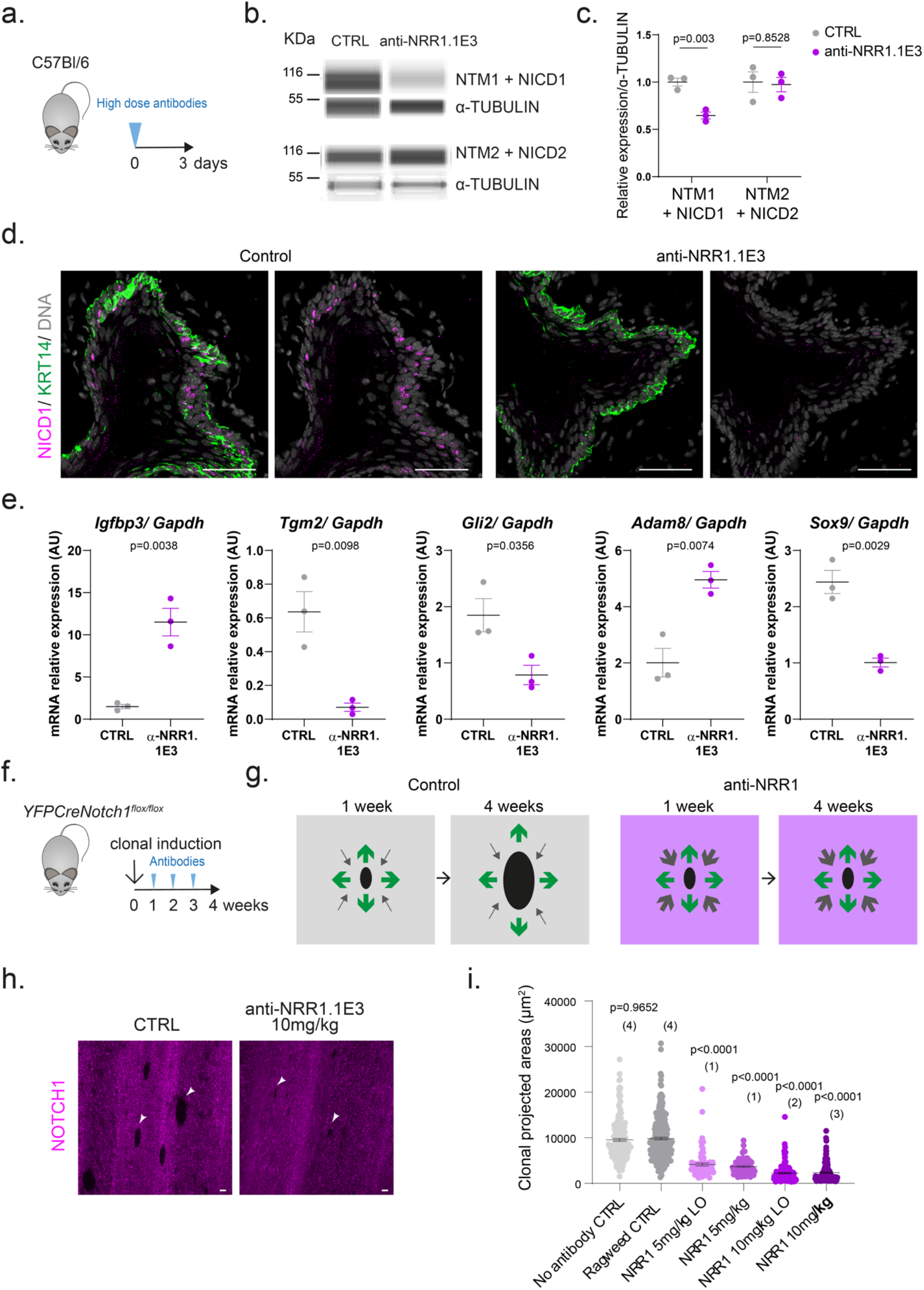
anti-NRR1 antibodies inhibit NOTCH1 signaling *in vivo*. **a.** Protocol for b-e. C57Bl/6 wild type mice were treated using neutralizing anti-NRR1.1E3 or anti- Ragweed control antibodies (CTRL) at high dose (25mg/kg) three days before tissue collection **b.** Immune Capillary Electrophoresis assay for the cleaved transmembrane and intracellular regions of NOTCH1 (NTM1 + NICD1, **Extended data Fig1a**) and NOTCH2 (NTM2 + NICD2), and α-TUBULIN proteins from esophageal epithelium of mice in **a. c.** Quantification of the relative expression of the cleaved transmembrane/intracellular regions of NOTCH1 and NOTCH2 (NTM + NICD) normalized on α- TUBULIN expression in the indicated conditions. Mean ± SEM, each dot represents a mouse, n=3 mice. Two-tailed unpaired Student’s t-test. **d**. Representative pictures of immunostaining for NICD1 (magenta), KRT14 (green) and nuclei (grey) in esophageal cryosections of mice treated with control or anti-NRR1.1E3 antibodies (n=3). Scale bars, 30 µm. **e.** Plots show RT-qPCR measurement of *Notch1* loss of function marker transcripts identified in the bulk RNAseq (**Supplementary table 5**) in control and anti-NRR1.1E3 treated samples. Mean ± SEM, each dot represents a mouse, n=3. Two-tailed unpaired Student’s t-test. **f.** Protocol for g-i. *YFPCreNotch1^flox/flox^* mice were induced at clonal density and antibody treatment started one week later. Mice were dosed with anti-NRR1.1E3 or control antibodies for 3 weeks before esophageal collection. **g.** Principle of assay shown in **f** to measure the efficacy of the long-term treatment. As expansion of clones is subject to the fitness of the neighbor cells ^19^, induced *Notch1^-/-^* cells treated with control antibody one week after induction have higher fitness (green arrows) than surrounding wild type cells (grey arrows) and clones expand to a bigger size until collection at 4 weeks. If the anti-NRR1.1E3 treatment blocks NOTCH1 signaling, the surrounding cells have a fitness identical to the induced *Notch1^-/-^cells* and the expansion of *Notch1^-/-^* clones will be halted. **h**. Representative images of NOTCH1 negative clones in esophageal epithelia treated with control or anti-NRR1.1E3 antibody at 10mg/kg and collected one week after the last dose. Images are representative of entire esophagus from 3 mice. Scale bars, 50µm. **i**. Quantification of area of clones negative for NOTCH1 staining. Mean ± SEM, each dot represents a clone. Number of mice analyzed per dose are indicated in brackets. One-way ANOVA; Tukey’s multiple comparisons test, adjusted p-values shown against Ragweed control condition. AU, arbitrary unit. LO, loading.

